# SACHET: seeded contour tracking measures Rab conversion kinetics on individual endosomes in live cells

**DOI:** 10.64898/2026.10.30.763220

**Authors:** Felix Margadant, Ann-Christin Borchers, Duarte Mateus, Anna Vik Rødseth, Vinodha Manovaseegaran, Xiaochun Xu, Michael P. Sheetz, Oddmund Bakke, Xian Hu

## Abstract

Membrane-labeled endosomes appear as hollow rings in a projection; they move, deform and exchange their coat, which population trackers — tools that detect every object of one kind and link the detections across frames — are not designed to follow. Here we introduce SACHET (Seeded Adaptive Contour & Hollow Endosome Tracking), a human-supervised Fiji tool that tracks one endosome at a time. On simulated ground truth it localizes the center to 10 nm, recovers transition time constants and lags without bias and holds the target among touching neighbors; comparisons with tuned TrackMate and Cellpose delineate where each is superior. In crowded spinning-disk projections, TrackMate followed one of seven marked Rab5-to-Rab7 conversions; supervised SACHET followed all 37. SACHET thus provides a stable per-endosome frame in which membrane morphology and protein exchange can be quantified: in five cell lines the conversion toggles like a digital switch within seconds to minutes, differing between lines.

## Introduction

Endosomes sort the material that cells take up by endocytosis and pass it on to recycling or degradation [1, 2]. Rab GTPases are key determinants of endosomal membrane identity [3–7]: Rab5a (hereafter Rab5) and Rab4 mark early endosomes, Rab11 recycling endosomes, and Rab7a (hereafter Rab7) and Rab9 late endosomes [8–12]. Maturation from early to late endosome is executed by a Rab conversion in which Rab5 is lost and Rab7 gained [13, 14]. The coupling is provided by the Rab7 guanine-nucleotide exchange factor (GEF) Mon1–Ccz1, a Rab5 effector that displaces the Rab5 GEF Rabex-5 [14–19], and by the Rab5 GTPase-activating protein (GAP) TBC1D18 [20]. Network models of this circuit suggest a “cut-out switch” mechanism analogous to a circuit breaker [21], and reconstituted Rab5 activation is sigmoidal and bistable [22]. Whether the conversion on a living endosome is gradual or toggles, how fast it is and how reproducible it is between endosomes has not been measured directly: a systematic literature search (Methods) found no study reporting Rab5-to-Rab7 conversion times for individual mammalian endosomes, the closest being manual tracking of 14 enlarged endosomes [23], peak-aligned traces on ionophore-enlarged endosomes [24], qualitative scoring [25], model-constraining trajectories [21] and automated kinetics of the earlier APPL1-to-EEA1 transition [26].

The knowledge gap is rooted in an intricate tracking problem. A maturing endosome is 0.5–2 µm across and, in an optical section, appears as a ring or loop with a dim lumen rather than as an extended blob. During maturation it moves, squeezes through the cytoplasm, fuses, extends tubules and — as its membrane-protein concentrations change (Rab5 falling, Rab7 rising) — changes its appearance in each channel independently. The tools that dominate trafficking analysis are population trackers, which detect every object that shares an image signature — a spot, a mask — and link the detections across frames. Single-particle trackers such as TrackMate [27, 28] detect diffraction-limited spots and link them across a whole cell; a hollow ring of variable radius is not the object they were built to detect, and the disappearance of the Rab5 signal ends a Rab5-based track exactly when the measurement ought to commence. Deep-learning segmentation networks such as Cellpose [29] and interactive classifiers [30] outline every object in every frame, but a mask carries no identity and must be linked afterwards. Active contours [31] model a ring naturally but need per-dataset tuning and have no defined recovery when lost. Earlier work therefore enlarged endosomes with Rab5 Q79L [13], invariant-chain-driven fusion [32] or nigericin [24] so that a fixed region of interest can be manually outlined, which changes the biology and restricts the sample to a few, round, stationary endosomes. Imaging is a second constraint: endosomes move in three dimensions, and recording z-stack volumes rapidly enough to capture a conversion within tens of seconds photobleaches the fluorescent signal before the conversion completes. We developed the micro-stepping extended focus (EF) imaging system [33] for this problem: it records the cell depth as one optical projection, about ten times faster and with substantially less bleaching than a stack; most of the cell-line recordings were acquired with EF.

SACHET rests on a simple premise: an object larger than the diffraction limit, seen only in its projection, can be localized well below the optical resolution of the microscope by tracking the centroid of the observed membrane. EF supplies such projections without depth distortion, and SACHET turns them into a frame of reference that moves with the endosome, in which its shape and membrane intensities are read frame by frame. Because the contour detection tolerates low signal, recordings can be fast and long at low illumination, which resolves the conflict between speed, precision and duration described above.

Here we describe SACHET, a seeded, human-supervised tracker for ImageJ/Fiji that follows a single endosome through a two-channel time series with an annular contour re-fitted in every frame (Fig. 1). It is a mid- to low-throughput, high-resolution tool for one vesicle at a time, not a replacement for population trackers. We describe the algorithm and its Fiji pipeline (Figs. 1–2), assess our accuracy claims on synthetic ground-truth datasets including deforming shapes and crowded fields, compare it with tuned TrackMate and Cellpose (Figs. 3–4; Table 1), test it on fast, crowded spinning-disk recordings where automatic tracking largely fails (Fig. 5), and apply it to the Rab5-to-Rab7 switch in five cell lines, to a 16-condition co-expression screen of its speed and to eight further membrane proteins placed relative to the switch (Figs. 6–8). All code is freely available (Code availability).

**Fig. 1:**
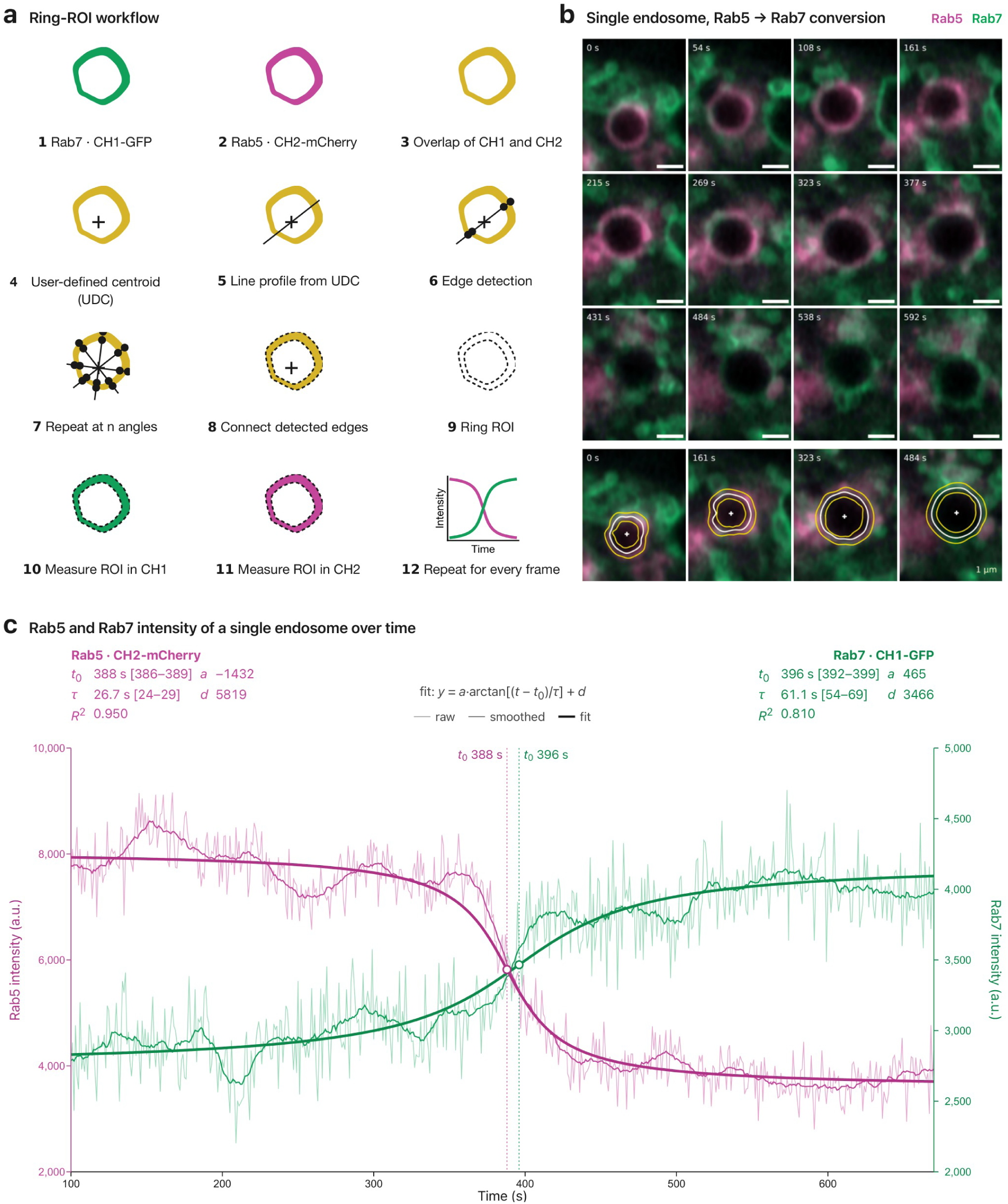
SACHET follows a single maturing endosome with an adaptive annular contour. **a**, Process steps as described in Results (Design of SACHET and its Fiji pipeline). **b**, Time-lapse of a maturing endosome in an MDCK cell co-expressing mCherry-Rab5 (magenta) and EGFP-Rab7a (green), single confocal plane, 1.35 s per frame (Supplementary Movie 1). Rows 1–3, merged channels at the indicated times; row 4, SACHET contour (white), the edges of its ±2 pixel measurement band (yellow) and the contour centroid (dot) at four time points. Scale bars, 1 µm. **c**, Mean membrane intensity of Rab5 (magenta, left axis) and Rab7 (green, right axis) in the SACHET ROI of the endosome in **b**: raw trace (thin), smoothed trace (medium) and arctangent fit I(t) = a·arctan((t − t0)/τ) + d (thick) with the fitted parameters and 95% confidence intervals; dotted lines and circles mark the transition midpoints (Rab5 t0 = 388 s, τ = 26.7 s; Rab7 t0 = 396 s, τ = 61.1 s; lag 8 s). n = 1 endosome; the population is shown in Supplementary Fig. 5.

**Fig. 2:**
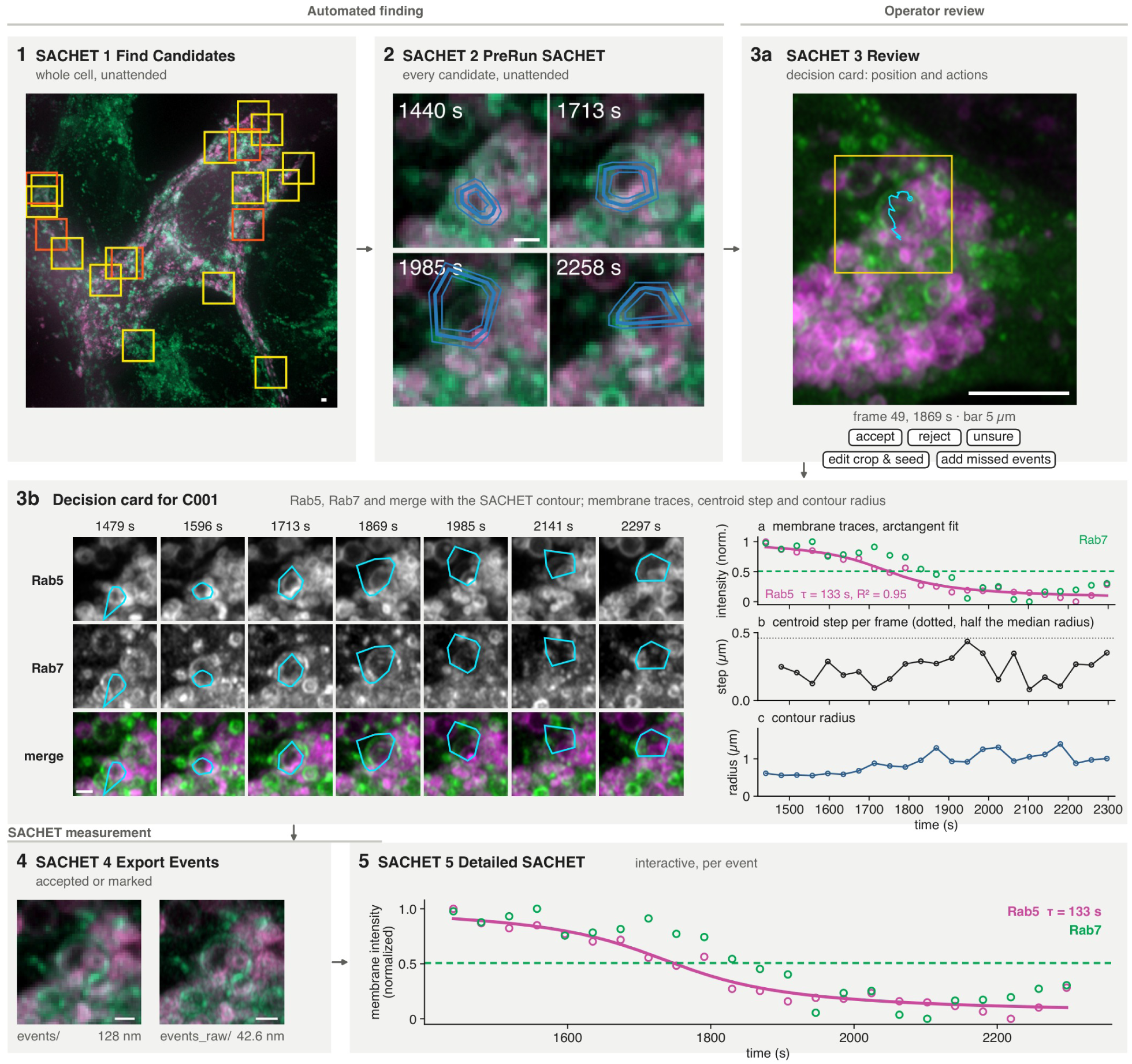
The SACHET Fiji pipeline from whole-cell movie to per-event kinetics. Steps 1–5 as described in Results (Design of SACHET and its Fiji pipeline). (1) Deconvolved widefield DeltaVision cell with the 19 crop boxes proposed by the candidate finder (yellow, candidate; orange, partial transition). (2) Pre-run of SACHET on Airyscan candidate C001 at 1,440, 1,713, 1,985 and 2,258 s; blue, SACHET contour and ±2 pixel measurement band. The pre-run tracked ≥ 80% of the frames for 2 of 19 DeltaVision and 2 of 7 Airyscan candidates. (3a) Review of the same candidate: its position in the cell at the reference frame (frame 49, 1,869 s; yellow, crop box; cyan, path of the SACHET contour centroid) and the actions available per candidate (buttons). (3b) Content of the decision card for C001: Rab5, Rab7 and merged channels at seven time points with the SACHET contour (cyan); a, membrane intensity of both channels (normalized) with the arctangent fit to Rab5 (τ = 133 s, R² = 0.95; dashed, mean Rab7); b, step of the contour centroid per frame (dotted, half the median contour radius, above which a frame is flagged); c, contour radius. (4) The same event exported on the analysis grid (events/, 128 nm) and at native resolution (events_raw/, 42.6 nm); all 7 exports reproduced the analysis image and the raw file exactly. (5) Unattended run of the interactive SACHET on C001: points, membrane intensity per frame (normalized); line, arctangent fit to Rab5 (τ = 133 s, R² = 0.95); Rab7 shows no rise on the contour (R² ≈ 0; dashed, its mean). Counts of the test on one DeltaVision and seven Airyscan cells are given in Results and the script runtimes in Supplementary Methods; review decisions in the test were scripted. Events marked by the operator where the automatic finder fails enter at step 4 (Fig. 5a, b). Rab5 magenta, Rab7 green throughout. Scale bars, 1 µm (1, 2, 3b, 4) and 5 µm (3a).

**Fig. 3:**
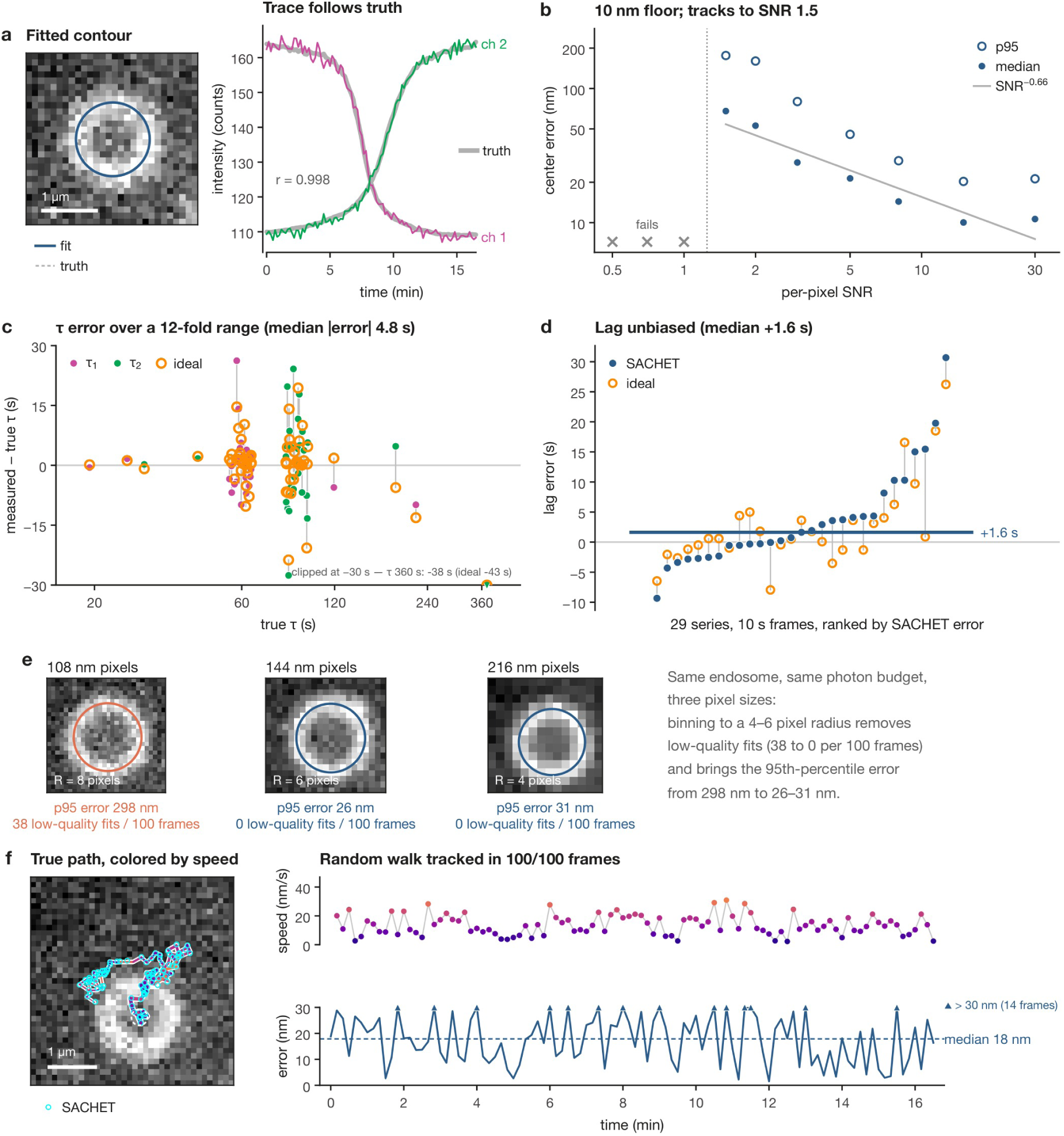
Validation of SACHET on synthetic ground truth. **a**, Left, one frame of a synthetic two-channel series (ring radius 6 pixels, SNR 8; channels summed) with the fitted SACHET contour (blue) and the true membrane (white dashed). Right, mean membrane intensity measured by SACHET in the falling (magenta) and rising (green) channel, overlaid on the noise-free ground truth (gray); r = 0.998. **b**, Center localization error (median, filled; 95th percentile, open) against per-pixel SNR for a 6 pixel ring; the line is a power-law fit to the medians (exponent −0.66). Crosses mark SNR values at which the contour could not be initialized. **c**, Error of the fitted transition time constant (measured − true τ) against true τ for both channels of every valid run (n = 60 fits); orange rings, the same error for the arctangent fit to the noise-free membrane intensity of the same series (ideal tracker), joined to the SACHET value by a gray line; horizontal line, zero error. One value lies below the axis (τ = 360 s: −38 s; ideal −43 s) and is drawn at its edge. **d**, Error of the lag between the channel-2 and channel-1 midpoints (measured minus true) for the 29 series at 10 s per frame (true lag 120 s), ranked by the SACHET error; filled, SACHET; orange rings, ideal tracker on the same series, joined to the SACHET value by a gray line; bar, median (+1.6 s). The series at one frame per second (true lag 13 s, recovered as 15.8 s) is given in the text. **e**, The same 0.86 µm endosome rendered with a constant photon budget at 108, 144 and 216 nm pixels (first frame with the fitted contour; R, ring radius in pixels); under each crop the 95th-percentile center error and the fit instability (low-quality fits per 100 frames). **f**, Random walk of 13 nm s⁻¹ median speed: true path colored by speed (tail thickens towards the final frame, dashed ring) with the SACHET centroid of every frame (cyan), and the speed and tracking error over time on a linear axis capped at 30 nm (triangles, the 14 frames with an error above 30 nm; dashed line, median 18 nm); 100/100 frames tracked. Scale bars, 1 µm. n = 42 synthetic series (33 valid runs); per-run values in Supplementary Fig. 1 and Source Data.

**Fig. 4:**
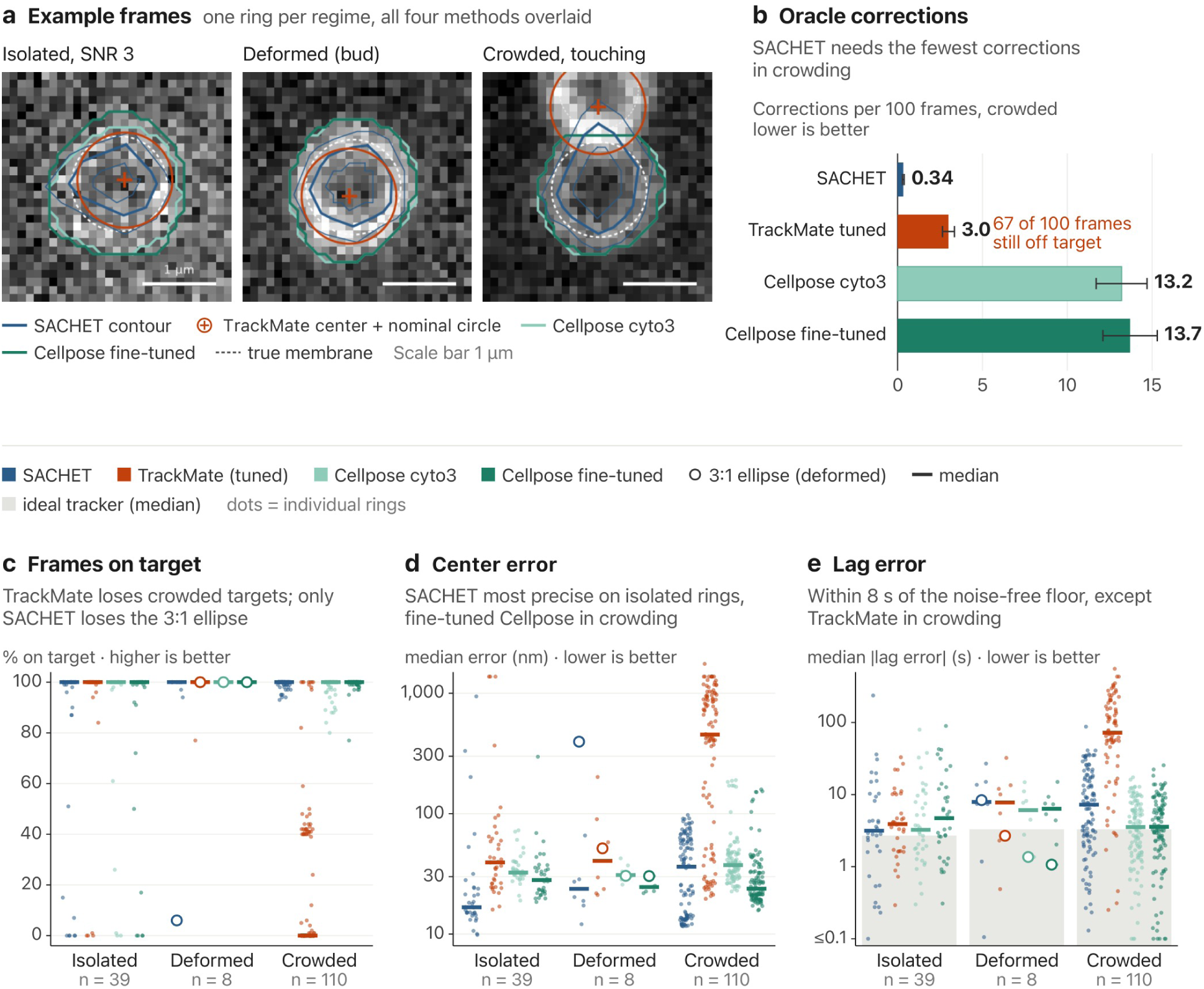
SACHET, TrackMate and Cellpose on isolated, deformed and crowded synthetic endosomes. Synthetic two-channel stacks with ground truth; every tracker was seeded at the frame-1 lumen center (SACHET round-ring values are the operator runs of Fig. 3, all else unattended). **a**, One frame per regime (summed channels): isolated ring at SNR 3 (frame 46), budding ring at SNR 10 (frame 50) and ring touching a Rab5-high neighbor (gap 0, frame 50). Blue, SACHET contour (thick) and its ±2 pixel measurement band (thin); orange-red cross and circle, TrackMate center with the nominal ring radius; light and dark teal, pretrained and fine-tuned Cellpose masks; white dashes, true membrane center-lines. Scale bars, 1 µm. **b**, Oracle corrections per 100 frames in crowding (mean ± 95% bootstrap CI over realizations): SACHET re-seeds, TrackMate link joins and cuts, Cellpose re-assign, merge and split events; 67 of 100 TrackMate frames had no on-target detection to correct. **c**, Frames on target: center within 2 pixels (216 nm) of the true center (membrane centroid for deformed shapes) and, in crowding, nearer the target than any other object; undetected frames count as off. **d**, Median center error per stack over detected frames (log scale). **e**, Absolute error of the fitted Rab5-to-Rab7 lag; gray bands, ideal tracker (noise-free image read in the true membrane band, median). Values below 0.1 s are drawn at 0.1 s; stacks without an acceptable fit (r² < 0.5) are omitted (fits: isolated 34, 35, 36, 35 of 39 for SACHET, TrackMate, Cellpose, fine-tuned; deformed 8 of 8; crowded 110, 79, 110, 110). In c–e dots are stacks and bars medians; open circles mark the 3:1 ellipse; values beyond the axis range are drawn at the top of the axis (center error > 1.3 µm, 11 TrackMate stacks; lag error > 430 s, 2 TrackMate stacks). Isolated, n = 39 (five stacks on which SACHET did not initialize count as 0%); deformed, n = 8 shapes at SNR 10; crowded, n = 110 (22 conditions × 5 realizations; isolated controls excluded). TrackMate configuration and Cellpose read-out as in Methods; all trackers, conditions and tuning results in Supplementary Figs. 3 and 4, Supplementary Data 1 and Supplementary Table 2. Tuning changed TrackMate’s crowded on-target fraction from 20.9 to 21.2%; the membrane-peak read-out reduced Cellpose’s median absolute lag error on isolated stacks from 32 and 27 s to 3.7 and 5.4 s (pretrained, fine-tuned); in the crowded fields TrackMate made 444 identity switches, Cellpose two and SACHET none. Source data are provided as a Source Data file.

**Fig. 5:**
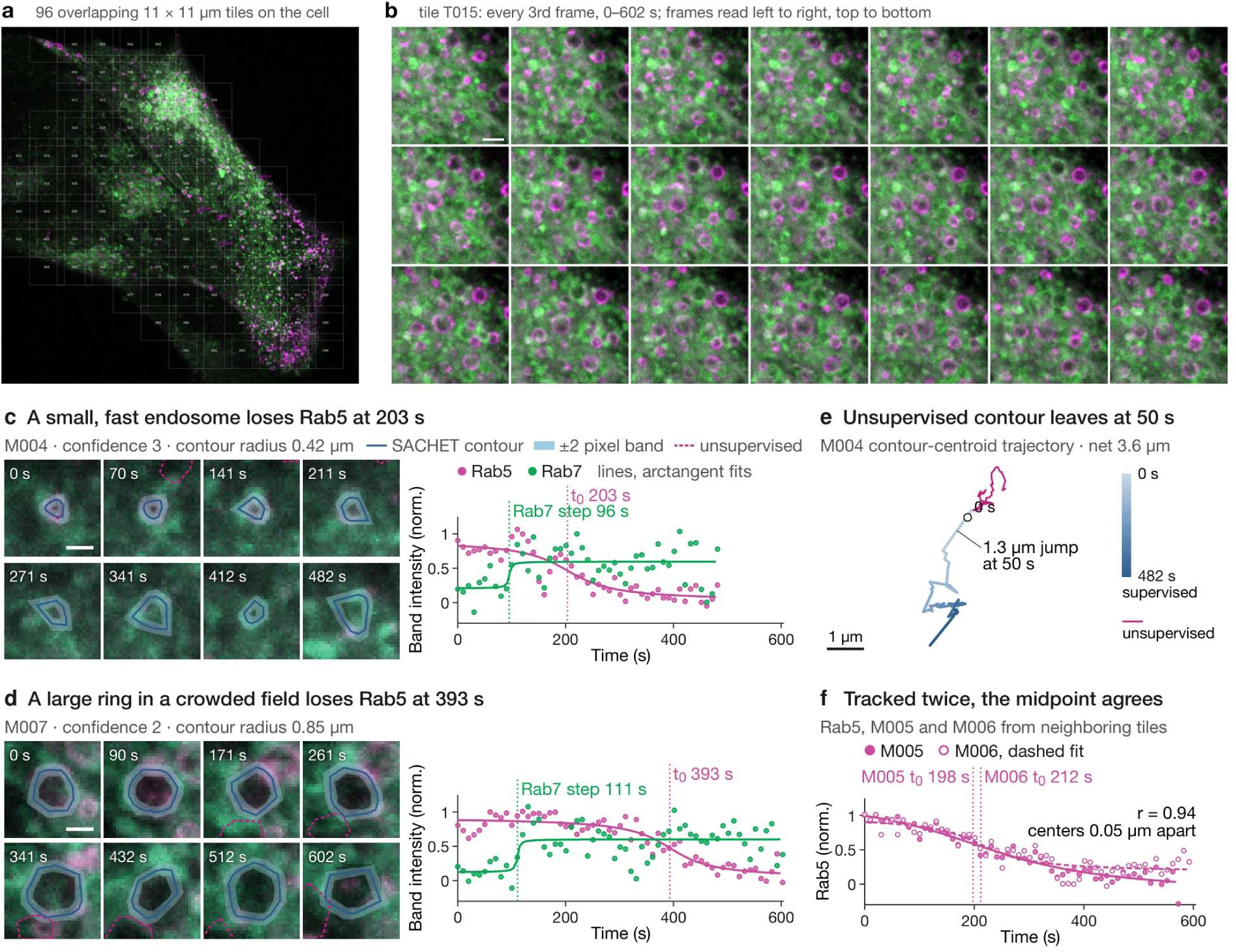
Supervised SACHET on fast, crowded endosomes in a spinning-disk recording. A different cell and imaging mode from Fig. 1: cell co-expressing EGFP-Rab5a (magenta) and mApple-Rab7a (green; colors denote the protein, not the fluorophore, throughout), 11-plane z-stacks every 10.04 s for 61 frames, 110 nm pixels, maximum-intensity projection; conversions marked by eye were tracked unsupervised (automatic re-seeding) and supervised (operator re-clicks; Supplementary Data 2). Position, radius and shape are those of the SACHET contour; intensities are the means in its ±2 pixel band. **a**, Systematic tiling of the cell into 96 overlapping 11 × 11 µm crops (20-pixel overlap), independent of any detector; numbers, tile identifiers. **b**, Contact sheet of tile T015 (every third frame, 0–602 s) as browsed by the operator, who records for each conversion the ring center, the frames at which the ring is clearly Rab5 and clearly Rab7, and a confidence score (1–3); marked events enter the pipeline at the export step (Fig. 2, step 4). Scale bar, 2 µm. **c**, Event M004 (confidence 3, contour radius 0.42 µm): merged channels at eight time points with the supervised SACHET contour (blue), its ±2 pixel measurement band (light blue) and the unsupervised contour (magenta, dashed); right, mean membrane intensity of Rab5 and Rab7 in the measurement band (normalized to the 2nd–98th percentile) with arctangent fits and fitted midpoints (dotted): Rab5 t0 = 203 ± 17 s, τ = 64 ± 31 s, R² = 0.80; Rab7 step at 96 ± 17 s, i.e. 107 s before the Rab5 midpoint (Δt0 = −107 s; τ at the 3 s bound), R² = 0.29. Scale bar, 1 µm. **d**, Event M007 (confidence 2, contour radius 0.85 µm, crowded field) as in c: Rab5 t0 = 393 ± 12 s, τ = 59 ± 19 s, R² = 0.88; Rab7 step at 111 ± 4 s (Δt0 = −282 s), R² = 0.43. **e**, Supervised contour-centroid trajectory of M004 colored by time (net 3.6 µm) with the unsupervised trajectory (magenta), which left the endosome at a 1.3 µm jump at 50 s. **f**, Reproducibility pair: Rab5 membrane intensity of the same endosome tracked from two neighboring tiles (M005, M006; normalized as in c, arctangent fits, M006 dashed); contour centroids 0.05 µm apart (median), traces correlated at 0.94, Rab5 t0 198 ± 26 and 212 ± 16 s, τ 187 ± 52 and 70 ± 28 s. n = 37 distinct endosomes from one cell; M002 (the track followed a neighboring ring 1.7 µm from the mark), M029 (operator: crowded, not accurate) and M030 (tracked M032’s endosome) are omitted; M006 (a repeat of M005) is shown only in f. Source data are provided as a Source Data file.

**Fig. 6:**
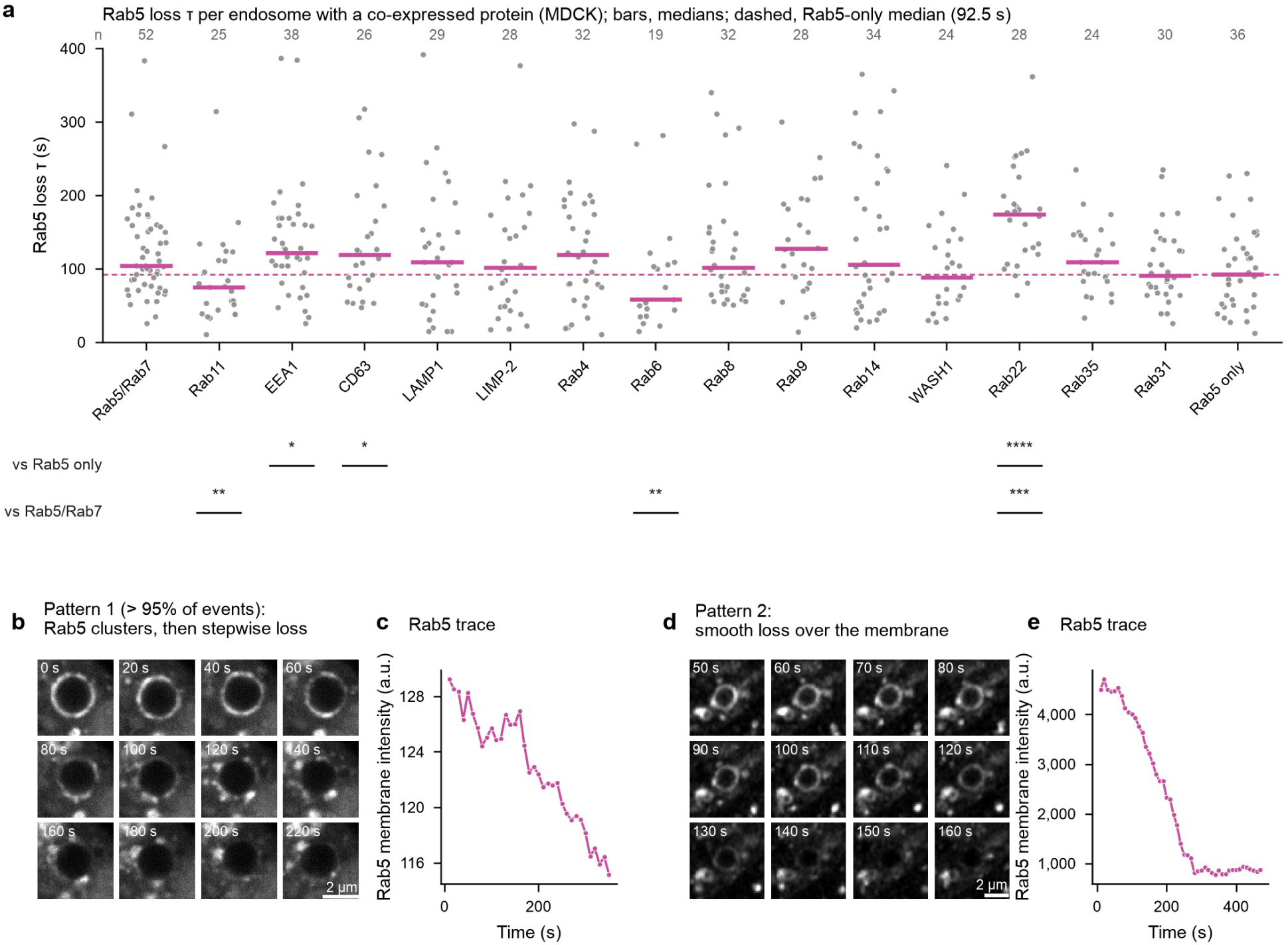
Rab5 loss kinetics under 15 co-expressed proteins, and the two forms of Rab5 loss. **a**, Rab5 loss time constant τ per endosome in MDCK cells expressing Rab5 alone (“Rab5 only”), Rab5 with Rab7 (“Rab5/Rab7”) or Rab5 with the indicated protein; each point is one endosome, horizontal bar the median of each group, dashed line the median of the Rab5-only group (92.5 s). Maturation events were scored by Rab5 loss; Rab7 gain followed Rab5 loss on every endosome in our dual-color recordings (Supplementary Fig. 5b) and in previous studies [13, 21, 23], so Rab5 loss alone is used where the second channel carried the co-expressed protein. n endosomes per group (above the panel): Rab7 52, Rab11 25, EEA1 38, CD63 26, LAMP1 29, LIMP-2 28, Rab4 32, Rab6 19, Rab8 32, Rab9 28, Rab14 34, WASH1 24, Rab22 28, Rab35 24, Rab31 30, Rab5 only 36; each group from ≥ 20 cells and ≥ 2 independent experiments. Kruskal–Wallis test followed by Dunn’s multiple-comparison test against the Rab5-only or the Rab5/Rab7 group as indicated (GraphPad Prism); two-sided; *p < 0.05, **p < 0.01, ***p < 0.001, ****p < 0.0001. **b, c**, Time-lapse (**b**) and SACHET Rab5 membrane-intensity trace (**c**) of an endosome showing pattern 1, the prevalent form (> 95% of events): Rab5 first concentrates into clusters on the membrane and then declines stepwise (Supplementary Movie 4). **d, e**, As **b, c** for an endosome showing pattern 2: progressive, smooth loss of Rab5 over the whole membrane (Supplementary Movie 5). MDCK cells, Rab5 channel shown in gray scale, 10 s per frame; time in seconds from the start of the recording; traces give the SACHET membrane intensity in arbitrary units. Scale bars, 2 µm. Source data are provided as a Source Data file.

**Fig. 7:**
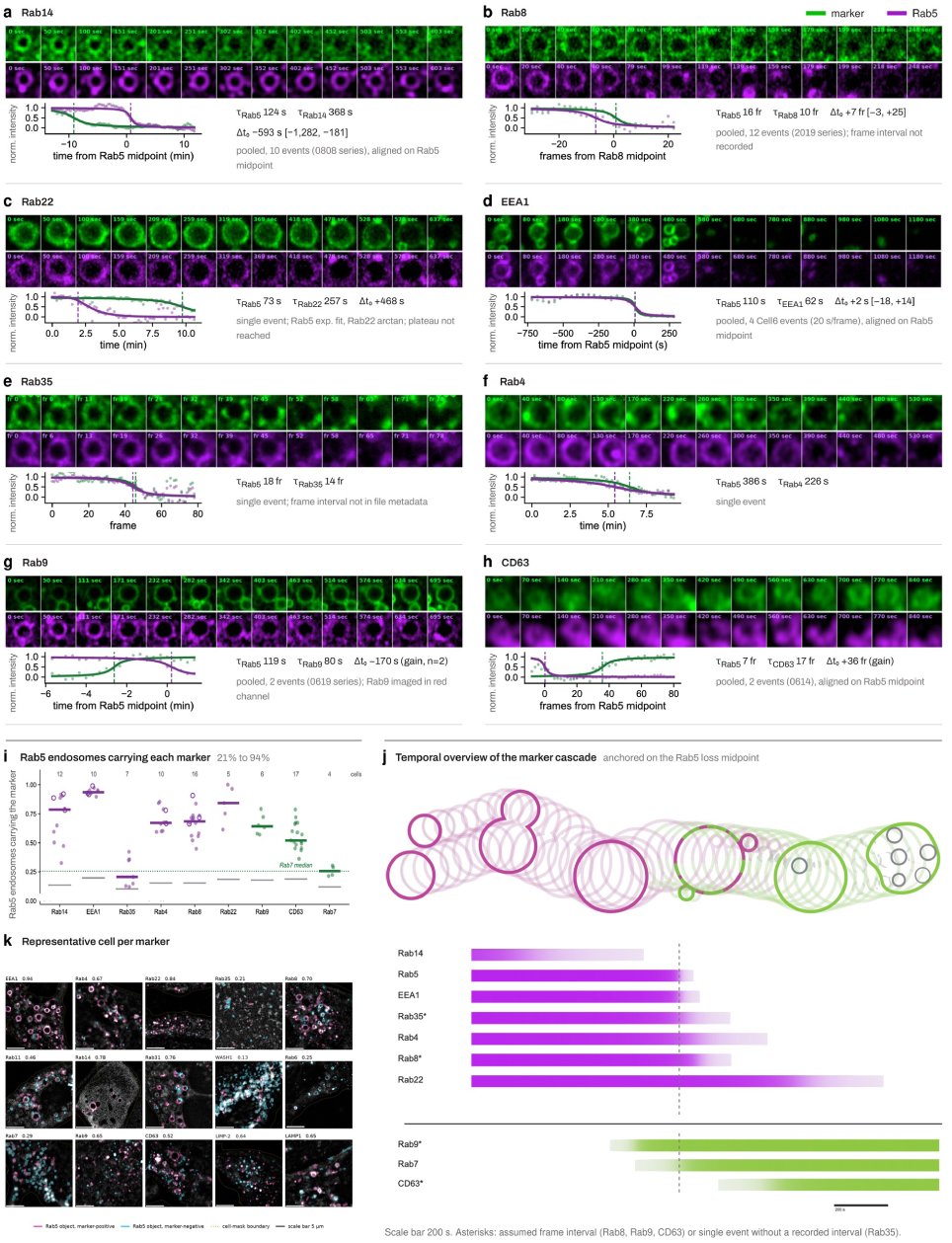
Placement of eight membrane proteins relative to the Rab5 loss midpoint on individual endosomes, and the fraction of Rab5 endosomes that carry each protein. **a–h,** Top, time-lapse of a maturing endosome (centered in each tile) in MDCK cells co-expressing Rab5 (magenta) with Rab14 (**a**), Rab8 (**b**), Rab22 (**c**), EEA1 (**d**), Rab35 (**e**), Rab4 (f), Rab9 (**g**) or CD63 (**h**) (green); time in seconds, or in frames where the interval was not recorded in the file metadata (Supplementary Table 4); each panel is displayed with its own linear intensity scaling, so intensities are not comparable between panels (Supplementary Movie 6). The montages are displayed without scale bars because they were assembled for an earlier version of the figure and the display scale of the crops was not recorded; for orientation, the endosomes shown are approximately 1–2 µm in diameter. Bottom, normalized SACHET membrane-intensity traces of the marker and Rab5 with fitted transitions (τ indicated) and the phase shift Δt0 of the marker midpoint relative to the Rab5 midpoint. For markers with several events the traces were aligned on the Rab5 (or Rab8) midpoint, pooled and fitted with the template (Methods): Rab14, 10 events; Rab8, 12; EEA1, 4; Rab4, 5; Rab9, 2; CD63, 2; the montage shows one representative event. Rab22 (c) and Rab35 (**e**) are single events; in **c** the Rab5 trace did not reach its plateau within the recording and was fitted with an exponential, the one exception to the acceptance criteria (Methods). Where n ≥ 4 the event-bootstrap 95% CI of Δt0 is given in brackets. Upon Rab22 co-expression Rab5-positive endosomes are enlarged (Fig. 8). **i**, Fraction of Rab5-positive endosomes scored as carrying each protein in single-time-point images of MDCK cells, for the eight proteins of **a–h** and Rab7 (proteins that lose intensity with Rab5 in the order of **j**, then those that gain it); one point per cell, horizontal bar the median; purple, intensity falls with Rab5; green, rises; gray bars, median randomized null (marker channel rotated, flipped or shifted, 20 iterations per image); open circles, cells flagged for possible spectral bleed-through; dotted line, Rab7 median (26%, n = 4 cells from two sessions); numbers above, cells per protein (1–5 imaging sessions; Supplementary Table 3). Rab7 cells of the invariant-chain-induced MDCK line (one session; induction enlarges endosomes, and 49–88% of its Rab7 puncta were Rab5-positive) and one low-signal Rab7 image were excluded at visual quality control; all 15 proteins are given in Supplementary Fig. 6a and Supplementary Table 3. **j**, Temporal overview anchored on the Rab5 loss midpoint (dashed line). Top, schematic of a maturing endosome on the same time axis (Rab5, magenta; Rab7, green; not to scale). Bottom, color stripes indicate the onset and duration of each protein’s transition (plateaus not shown); purple, proteins in phase with Rab5 loss; green, proteins in phase with Rab7 gain; scale bar, 200 s. Asterisks mark entries resting on an assumed frame interval (Rab8, Rab9, CD63) or on a single event without a recorded interval (Rab35). Only Rab14 and EEA1 are supported by several events with a recorded frame interval; Rab22 and Rab4 rest on single fully captured events; Rab9 and CD63 on two events each. Per-marker event counts and confidence classes are in Supplementary Table 4. **k**, One representative cell for each of the 15 proteins scored (the cell closest to its protein’s median): Rab5 channel with every detected Rab5 endosome outlined, magenta where the protein was scored present and cyan where absent; dotted line, cell mask; the number is that cell’s marker-positive fraction. Scale bars, 5 µm. **i**, **k**, n = 195 cells (191 with ≥ 10 detected Rab5 endosomes, used here and in Supplementary Fig. 6) over 15 proteins; stricter cutoffs, Manders coefficients and the relation to Δt0 are in Supplementary Fig. 6. Source data are provided as a Source Data file.

**Fig. 8:**
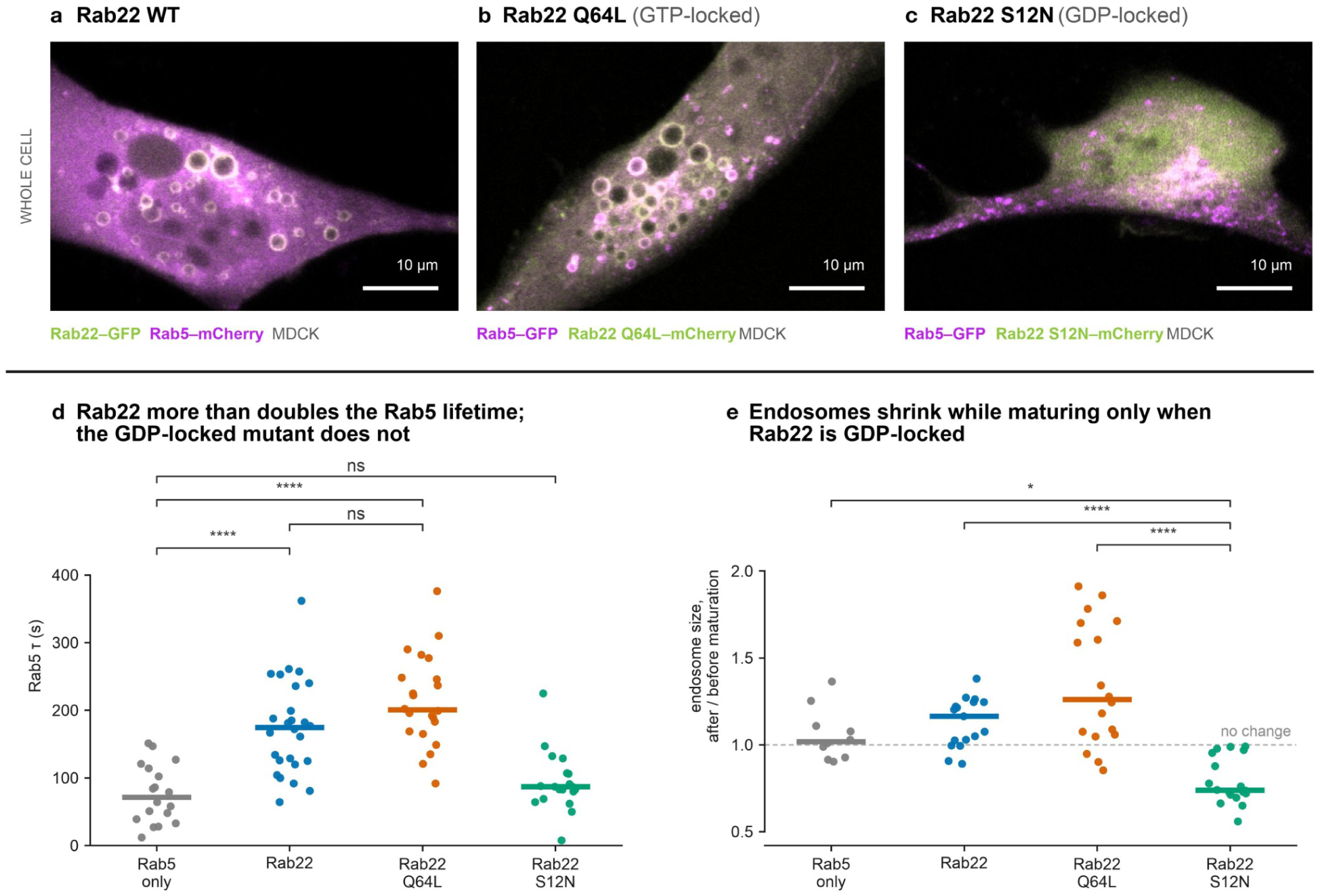
Rab22 activity slows Rab5 loss and enlarges Rab5-positive endosomes in MDCK cells. **a–c,** Live-cell confocal images of MDCK cells co-expressing Rab5 (magenta) with wild-type Rab22 (**a**), GTP-locked Rab22 Q64L (**b**) or GDP-locked Rab22 S12N (**c**) (green); one representative cell per condition, mean projection over the z-stack; all fields 54 × 36 µm at identical magnification; fluorophore assignments below each panel; linear intensity scaling per channel and panel (Supplementary Movie 7). Wild-type and Q64L Rab22 coat enlarged Rab5-positive endosomes, visible as rings up to 5.8 µm across; with S12N no ring is resolved. Scale bars, 10 µm. **d**, Rab5 loss τ per endosome for cells expressing Rab5 alone (n = 18) or with Rab22 (n = 26), Rab22 Q64L (n = 22) or Rab22 S12N (n = 19); medians 71.5, 174.5, 200.5 and 87.0 s, respectively. **e**, Endosome diameter after maturation relative to before, for the endosomes in **d** whose diameter could be measured at both times (n = 10, 17, 18, 18); dashed line, no change; medians 1.02, 1.16, 1.26 and 0.74. **d, e**, Each point is one endosome, pooled from two independent experiments per group; horizontal bars, medians. Kruskal–Wallis across the four groups (**d**: H = 45.1, p = 8.6 × 10⁻¹⁰; **e**: H = 33.4, p = 2.7 × 10⁻⁷) followed by Dunn’s test over all six pairs with Holm correction; brackets show the comparisons drawn; two-sided; ns p > 0.05, *p < 0.05, ****p < 0.0001. The Rab5-only group here is a separate experiment from the Rab5-only group of Fig. 6a and from the Rab5/Rab7 recordings of Supplementary Fig. 5, and serves as the within-experiment control. Diameters in **e** are ratios and unaffected by the +1 pixel calibration.

**Table 1:**
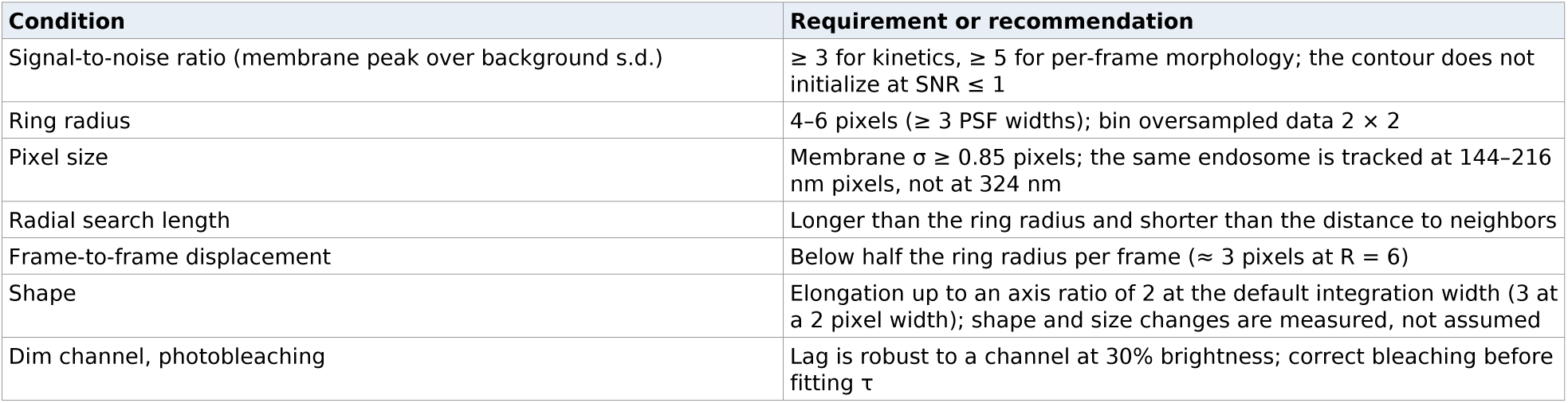
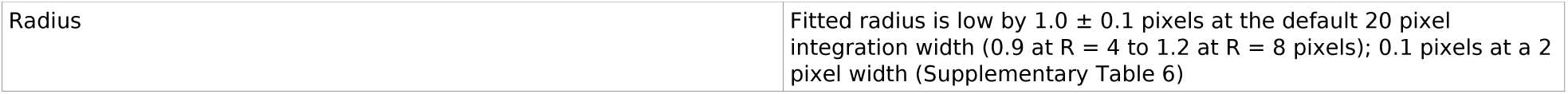
Operating window of SACHET on synthetic ground truth. Requirements for reliable tracking and the recommendation derived from each test; the measured values behind each row are in Supplementary Table 1, Fig. 3 and Supplementary Fig. 1; all rows refer to the default profile integration width of 20 pixels (Methods; Supplementary Table 6 for 2 pixels).

## Results

### Design of SACHET and its Fiji pipeline

The algorithm core is automated but requires supervision. Its input is a two-channel time series and a single point placed by the operator inside the lumen of the endosome of interest; that selection seeds the track, and the contour is fitted to both channels, so that the ring stays defined while one coat is lost and the other gained. The process has twelve steps (Fig. 1a) in four stages.

#### One membrane image (steps 1–3)

The two channels are equalized to a common dynamic range and combined into a single intensity image, a weighted sum of the two. All geometry is derived from this image, so the contour does not depend on the fluorophore brightness, the expression levels, or the dominant coat.

#### From seed to contour (steps 4–8)

From the user-defined centroid (4) a line profile is taken across the membrane (5) and the latter is located by edge detection with sub-pixel oversampling (6). This is repeated at n angles (20 at 18° by default; 7) and the detected edges are connected into a closed contour (8). The contour is a star domain — every ray from the seed crosses the membrane once — so deformations and protrusions are admissible as long as the seed lies inside the endosome.

#### The region of interest (ROI; step 9)

The connected endpoints form a closed-loop strip that contains the membrane projection with a 2-pixel margin on either side; it defines where the measurements are taken.

#### Measurement and propagation (steps 10–12)

The ring ROI is applied to each original channel (10, 11) and the mean intensity in the band is recorded together with the centroid, area, perimeter, circularity and median radius. The centroid seeds the next frame and the cycle repeats (12), without the need for a motion model.

The contour follows the endosome as it moves, deforms and extends a protrusion (Fig. 1b), and the Rab5 membrane intensity falls while Rab7 rises along sigmoidal curves; an arctangent fit to each trace yields its midpoint t0 and time constant τ (Fig. 1c).

#### Supervision

Where the contour is lost — typically after a displacement of more than half the ring radius or a fusion — the operator re-seeds at the failing frame and tracking resumes from there (Supplementary Movie 2). This cost is reported with every measurement (Results).

#### Implementation and practical use

Finding the events is addressed by a five-script Fiji pipeline (Plugins ▸ SACHET; Supplementary Software 2) in three stages (Fig. 2). (1) TrackMate tracks every endosome in the cell, and arctangent fits with rule-based filters propose Rab5-to-Rab7 conversion candidates. (2) SACHET then runs on the candidates. (3) The tracks are presented to the operator for an accept/reject/unsure/seed/edit crop/add decision. Where the automatic finder fails, the operator marks events on tile contact sheets and they enter the pipeline at step 4 (Fig. 5a, b; Results). (4) The tracks are exported and (5) the measurements and fits are performed and presented.

### Validation on synthetic ground truth: precision, completion and kinetics

We quantified SACHET on 42 synthetic two-channel series (Methods; Fig. 3; Supplementary Table 1; Supplementary Fig. 1). Each contains the projection of one membrane-labeled endosome, a loop with a falling and a rising channel of arctangent kinetics, rendered with noise and moving background puncta at the pixel size (108 nm) and frame intervals (10 s or 1 s) of our recordings; they vary signal-to-noise ratio (SNR), radius, pixel size, transition speed, motion, shape, photobleaching and channel brightness. Kinetics were compared with the truth and with an “ideal tracker”, the fit to the noise-free membrane intensity, separating tracking error from fitting error (Methods).

For a 6 pixel (0.65 µm) ring at an SNR of 8 the tracked center lay 14 nm (median) from the true center and the membrane intensity followed the noise-free trace with r = 0.998 (Fig. 3a). Localization error fell with SNR as a power law to a floor near 10 nm (0.1 pixel) at SNR ≥ 15 (Fig. 3b); all frames were tracked without re-seeding down to SNR 1.5; the contour could not be initialized at SNR ≤ 1. τ was recovered over a 12-fold range (true τ 20–240 s, 60 channel fits; median absolute error 4.8 s, 6%; Fig. 3c); SACHET and the ideal tracker deviated from truth alike (−0.4 ± 5.0 s at τ = 60 s, n = 26), so the scatter arises from noise and frame interval, not from tracking. The lag between the channel midpoints was recovered without bias (median error +1.6 s, interquartile range −0.5 to +4.3 s, 29 series at 10 s per frame; Fig. 3d), and a 13 s lag in the series recorded at one frame per second was recovered as 15.8 s (ideal tracker 15.1 s).

Tracking quality depended on the pixel size at which the endosome is sampled rather than on the photon budget: rings of 4–6 pixel radius gave ≤ 1% low-quality fits, 8 pixel rings 27–38% (0% at a 2 pixel width; Supplementary Fig. 2), and the same endosome rendered with the same photon budget at 144 or 216 nm pixels was tracked with 13–15 nm precision (Fig. 3e); rings below three optical point-spread functions (PSF) in radius have no resolvable lumen. Displacements of 1.2 pixels per frame were followed with 18 nm precision; at 3 pixels per frame, half the radius, the track was lost in 49 of 100 frames (Fig. 3f). Shape changes, distractors, a dim second channel and photobleaching were tolerated; the fitted radius was low by 1.04 ± 0.10 pixels, an effect of the integration width (Methods; Supplementary Fig. 2). Table 1 gives the operating window.

### Deformation, signal-to-noise ratio and size

Because endosomes elongate, bud, squeeze and carry uneven coats, we extended the ground truth to ten deformed shapes (Supplementary Fig. 3a) and ran SACHET unattended against TrackMate and generic trackers (Methods; Supplementary Fig. 3b). On the six stacks with moderate deformation (bean, amoeboid shapes, moving bud, polarized coat) SACHET followed all frames with 22 nm median center error, against 46 nm for TrackMate, 35 nm for trackpy [34] and 64 nm for the weighted centroid; the polarized coat displaced every point detector by 1–2 pixels (TrackMate, for example, by 208 nm) but the contour centroid by only 16 nm. A drifting ring squeezed to a 2:1 peanut was followed through its waist (2 frames off, 19 nm, lag error 1 s), whereas the fixed-radius band of the point trackers added 20–30 s of lag error during the constriction; at SNR 3 the squeeze became unstable (21 frames off). Elongation is the limit: radial profiles under-fill the ends of an ellipse, so the contour enclosed 0.74 of the true area at axis ratio 2 (4 frames off, 70 nm) and 0.54–0.61 at axis ratio 3, where the contour collapsed onto one end in 91 frames and was off target in 78–98 frames at SNR 8–30, unlike TrackMate and the centroid (Source Data). At a 2 pixel integration width the 3:1 ellipse was followed in every frame (19 nm; Supplementary Table 6), so the limit is the tangential averaging of the default width, not noise; axis ratio 2 is within range at both.

### Crowding and identity

To test the contour in crowding — neighboring endosomes whose projections touch or overlap — we added neighbors at controlled gaps to 110 synthetic stacks (Methods; Supplementary Fig. 4a). Unattended, SACHET was on target in 99.5% of frames and never switched identity.

Detection-and-linking failed when blurred membranes touched: TrackMate and trackpy were on target in 21% and 13% of frames, Otsu thresholding [35] merged target and neighbor (25%), and watershed separated touching rings but not puncta clusters or dense fields (3–12%). Pretrained Cellpose, its masks linked between frames by proximity, matched SACHET (98.8%; fine-tuned 99.5%), so holding the target among touching neighbors is not unique to the contour. Staying on target did not abolish leakage: beside a Rab5-high ring the neighbor entered SACHET’s trace with a coefficient of 0.25, like Cellpose (0.20) and watershed (0.28; ideal 0.11), whereas the point trackers mostly measured the neighbor (0.68–0.79). Beside a neighbor of the wrong kinetics the median absolute lag error was 16 s for SACHET, 18 and 14 s for Cellpose (pretrained and fine-tuned), 36–51 s for threshold and watershed and ≥ 150 s, or no fit, for the point trackers (ideal 5 s). With oracle correction SACHET needed 0.34 re-seeds per 100 frames, Cellpose 13, watershed 23 and threshold 84; TrackMate and trackpy needed three edits but had no on-target detection in two-thirds of frames.

### Comparison with TrackMate and Cellpose

We tuned TrackMate and Cellpose (Methods) before scoring all three on isolated round rings (39 stacks), deformed shapes (8) and crowded fields (110) (Fig. 4a). Tuning changed TrackMate little (Supplementary Data 1); Cellpose’s intensity was read at the membrane peak inside its mask, which cut its lag error on isolated stacks from 27–32 s to 4–5 s (Supplementary Table 2). On isolated rings all trackers held the median stack (Fig. 4c); SACHET had the lowest center error (17 nm versus 28–39 nm; Fig. 4d) but failed on five stacks (SNR ≤ 1, R = 3 pixels, 324 nm pixels) and was unstable on 20 pixel rings. Cellpose lost most frames at SNR 1.5–2. On deformed shapes SACHET lost the 3:1 ellipse, which every other tracker followed, and had the lowest center error on the bud and the polarized coat (12 and 17 nm versus ≥ 24 nm) but a larger lag error than Cellpose on the squeeze and the polarized coat. In crowding (above) Cellpose’s lag error (3.5–3.6 s) sat at the noise-free floor (3.4 s), half of SACHET’s (7.2 s; Fig. 4e), and fine-tuned Cellpose had the lowest center error (24 versus 36 nm). SACHET’s advantage in crowding lies in the correction cost: an oracle operator made 0.34 corrections per 100 frames with SACHET, 3.0 with TrackMate (67 frames per 100 still unrecoverable) and 13–14 with Cellpose, where most corrections repaired masks that had split or merged with a neighbor (Fig. 4b). SACHET thus matches a deep-learning segmentation network in crowding without training data and is the most precise tracker on isolated and moderately deformed rings, though not the most accurate in every regime.

### Real-data validation: supervised tracking of fast, crowded endosomes

We then tested SACHET on a spinning-disk z-stack of a cell co-expressing EGFP-Rab5a and mApple-Rab7a (11 planes every 10.04 s for 61 frames, 110 nm pixels, maximum-intensity projection; Methods), a difficult case: ring-shaped endosomes of 0.4–0.9 µm radius crowd the perinuclear region and move 0.1–0.4 µm per frame (median) with jumps of up to 2.3 µm, four ring radii. Lacking ground truth, the SACHET operator marked Rab5-to-Rab7 conversions by eye on contact sheets of 96 overlapping 11 × 11 µm tiles (Fig. 5a, b) with a confidence score (1–3), yielding 48 distinct endosomes, of which 37 are reported and 11 excluded (Supplementary Data 2).

Automatic tracking largely failed here. TrackMate, with ring-template detection and tuned linking, produced a track covering the conversion for one of the first seven marked events (M005) and none for the other six. Unsupervised SACHET with automatic re-seeding placed its contour centroid more than 0.5 µm away from the supervised contour centroid in a median 64% of frames (0–100%, 35 endosomes) and held the target throughout on seven slow, isolated ones; generic preprocessing did not change this (24 events; Source Data). Cellpose on the same crops held the target in a median 52.5% of frames (cyto3; fine-tuned 44.3%; unsupervised SACHET 36%) and needed a median 36 (fine-tuned 43) oracle corrections per 100 frames against the operator’s 19 re-clicks (Source Data).

Supervised SACHET followed all 37 endosomes (Fig. 5; Supplementary Data 2). The operator re-clicked the ring whenever the contour was lost: a median of 10 re-clicks per event (range 0–50), or 17.6 re-clicks per 100 frames (median; range 0–89; Supplementary Data 2). Rab5 loss was fitted in 11 of 37 endosomes (R² 0.74–0.93). Event M004 (radius 0.42 µm; Fig. 5c, e; Supplementary Movie 3) lost Rab5 with t0 = 203 ± 17 s and τ = 64 ± 31 s (R² 0.80) while traveling 3.6 µm net with steps of up to 1.3 µm; the unsupervised contour lost the endosome at the first such jump (50 s) and stayed 1.4–4.6 µm away. M007 (radius 0.85 µm), a large ring in a crowded field, lost Rab5 with t0 = 393 ± 12 s and τ = 59 ± 19 s (R² 0.88; Fig. 5d). The same endosome tracked twice from neighboring crops (M005, M006) gave centers 0.05 µm apart (median), Rab5 traces correlating at 0.94 and midpoints of 198 ± 26 and 212 ± 16 s, although τ differed (187 ± 52 versus 70 ± 28 s), suggesting that the midpoint is the more reproducible quantity (Fig. 5f).

The Rab7 gain was small (+6 to +19% where the fit was bounded), as mApple-Rab7a on these endosomes is barely brighter than the cytosolic pool. Its fitted rise preceded the Rab5 midpoint in three endosomes (M004, M007, M015) and followed it in two (M005, M011), counting the five endosomes with a resolved Rab5 midpoint and a bounded Rab7 fit; with five events from one cell and weak Rab7 fits (R² 0.29–0.73; τ at the 3 s bound in M004 and M007), this ordering — to our knowledge not previously reported — is weakly supported.

### Automatic candidate finding on slower recordings

The automatic finder (Fig. 2, steps 1–3) is meant for recordings slower and sparser than the spinning-disk movie of Fig. 5. On a cell imaged by deconvolved widefield (DeltaVision) TrackMate produced 1,284 tracks and the rules proposed 19 candidates, none of which was a resolved ring. On seven Airyscan cells, ring-mode detection produced 1,031 tracks and 7 candidates; five of them appear as puncta and none shows a clean conversion on its card. The finder thus runs end to end and shortlists candidates, but found no true conversion in these recordings. On the fast, crowded spinning-disk movie of Fig. 5, however, the finder returned none of the marked events, and the operator-marked entry point (Fig. 5a, b) was used instead.

### Application 1: the Rab5-to-Rab7 conversion toggles like a digital switch within seconds to minutes and differs between cell lines

A maturation event is defined as the loss of Rab5 from an individual endosome, accompanied where recorded by the gain of Rab7; its speed is the Rab5 τ. Cells expressing tagged Rab5 and Rab7 were imaged as z-stacks every 10 s for about 20 min (Methods), and stochastic events were seeded and tracked with SACHET.

We measured the switch in five cell lines: epithelial MDCK, carcinoma HeLa, primary HFF fibroblasts and two melanoma lines available in the laboratory and known to differ in behavior, A2058 and MelJuSo. The Rab5 τ of individual endosomes spread over more than an order of magnitude within every line, in right-skewed rather than Gaussian distributions (Supplementary Fig. 5a; MDCK: skewness 1.8 for Rab5 and 1.9 for Rab7 τ, Shapiro–Wilk p < 0.001 for both). Medians were 104 s in MDCK (n = 45), 62.5 s in HeLa (18), 170 s in HFF (10), 64 s in A2058 (25) and 125 s in MelJuSo (19); the 117 events ranged from 14 to 393 s, with 90% between 28 and 231 s. The τ distributions differed between lines (Kruskal–Wallis H = 27.6, p = 1.5 × 10⁻⁵): HeLa and A2058 were faster than MDCK and MelJuSo, and HFF was slower than HeLa and A2058 (Dunn–Holm p ≤ 0.02); the other pairs did not differ. The Rab7 gain τ ranked the lines the same way, with medians from 64 s (MDCK, n = 42) to 146 s (MelJuSo, n = 19); H = 24.6, p = 6.1 × 10⁻⁵ (Supplementary Fig. 5a).

In every line the Rab7 midpoint lagged the Rab5 midpoint, but by a variable amount (Supplementary Fig. 5b): median 84 s in MDCK (IQR 26–201 s; mean 123 s; positive in 38 of 45 events), 155 s in HeLa (12 of 16), 200 s in HFF (10 of 10) and 75 s in A2058 (20 of 20), with no detectable difference between lines (H = 5.8, p = 0.12); no endosome lost Rab5 without gaining Rab7. Pooled over 91 events the distribution was right-skewed with a tail to 844 s; in 11 the Rab7 fit placed its midpoint before that of Rab5. Rab7 timing is therefore not fixed by Rab5 loss alone. Every accepted trace was better described by the arctangent than by a monoexponential (Methods): the loss toggles like a digital switch rather than decaying as a first-order process. In more than 95% of events Rab5 first clustered on the membrane and then declined stepwise (pattern 1); in the remainder it declined smoothly (pattern 2) (Fig. 6b–e; Supplementary Movies 4, 5).

### Application 2: a co-expression screen of Rab5 loss kinetics

A spread this wide and skewed is compatible with several populations of endosomes maturing at different rates, for instance differing in the other proteins they carry. We therefore co-expressed Rab5 in MDCK cells with Rab7 or with each of 14 further proteins reported on early or maturing endosomes and measured the Rab5 τ on 485 endosomes (Fig. 6a; Supplementary Table 3; n = 19–52 per group). Most proteins left the kinetics unchanged (medians 88–128 s versus 92.5 s for Rab5 alone); Rab11 and Rab6 accelerated the loss (75 and 58 s), EEA1 and CD63 slowed it slightly (122 and 119 s) and Rab22 slowed it most (174.5 s). The spread persisted within every group. The screen also shows the scale at which SACHET can be used: 485 single-endosome time constants across 16 conditions.

### Other membrane proteins relative to the switch

Because both channels are measured on the same ROI, any protein co-expressed with Rab5 can be placed relative to the Rab5 midpoint by the midpoint difference Δt0. In the recordings of the screen, eight proteins showed a fittable transition on the same endosome as Rab5 (Fig. 7a–h; Supplementary Movie 6): EEA1, Rab8, Rab14, Rab22, Rab4 and Rab35 lose intensity and Rab9 and CD63 gain it; Rab11 forms transient blinking domains, and the rest showed no fittable change. Partial transitions were fitted with a template, and proteins with several events were aligned on the Rab5 midpoint, pooled and fitted with bootstrap confidence intervals (Methods; Supplementary Table 4). EEA1, a direct Rab5 effector [36], departed 2 s after Rab5 (95% CI −18 to +14 s; four pooled events; Fig. 7d). Rab8 departed with or shortly after Rab5 (Δt0 = +7 frames, CI −3 to +25; 12 events; Fig. 7b). Rab14 left well before Rab5 (−593 s, CI −1,282 to −181 s, ten events; individual fits −447 to −532 s; Fig. 7a). Rab4 departed after Rab5 (+18 frames, CI +8 to +45; five events; Fig. 7f), Rab22 468 s after (single event; Fig. 7c) and Rab35 in one of 31 recordings (+2 frames; Fig. 7e). Rab9 gain reached its midpoint about 170 s before the Rab5 midpoint (two events; Fig. 7g) and CD63 gain about 6 min after the Rab5 midpoint (36 frames; two events; Fig. 7h). Fig. 7j assembles these placements on a common axis.

Because these placements describe only endosomes carrying both proteins, we also scored in single-time-point images how often each protein is on a Rab5-positive endosome at all (Fig. 7i, k; Supplementary Fig. 6; Supplementary Table 3; 195 cells): EEA1 was on 94% of Rab5 endosomes, Rab22 on 84%, Rab14 on 79%, Rab8, Rab4 and Rab9 on 64–68%, CD63 on 52% and Rab35 on 21%, all above chance (10–20%), and Rab7 on 26%, as expected for a protein present only around the switch. Rab14 (leaving 590 s before Rab5, yet on 79%) and Rab9 (gained before Rab5 loss, yet on 64%) cannot be explained by brief residence windows, so these transitions change the amount of a protein already present: the endosome carries a mixed coat throughout.

Rab22 was examined further: in a separate experiment with its own Rab5-only control, wild-type and GTP-locked (Q64L) Rab22 increased τ more than twofold (medians 174.5 and 200.5 s versus 71.5 s for Rab5 alone; n = 26, 22, 18), whereas GDP-locked Rab22 S12N did not (87.0 s; n = 19), and both active forms enlarged Rab5 endosomes (Fig. 8; Supplementary Movie 7).

## Discussion

SACHET sits between the two classes of tool that dominate live-cell trafficking analysis, and the comparison in Fig. 4 defines that position rather than ranking the tools. TrackMate [27, 28] and Cellpose [29] focus on populations of signals and, on round, well-separated endosomes, are the superior choice, and with a membrane-centered read-out Cellpose recovered the kinetics as well as or better than SACHET on deformed and crowded synthetic rings. SACHET instead follows the projection of one chosen endosome as a ring with a lumen, re-fitting an annular contour in every frame on the sum of both channels so that the coat exchange does not interrupt the track. SACHET was developed alongside EF [33], which keeps a moving endosome in a fast, low-bleach projection of the cell depth, at the price that endosomes at different depths overlap — the crowded regime of Fig. 4.

The design has costs that define its scope. SACHET is supervised: on the fast, crowded spinning-disk recording of Fig. 5 the operator re-clicked the ring a median 18 times per 100 frames. That effort yielded a measurement the automatic alternatives did not deliver, but it bounds throughput. The real-data validation rests on one cell, one movie and a dim mApple-Rab7a channel, and its correction counts are not comparable with the oracle corrections of the synthetic benchmark, which omits focal drift, out-of-focus neighbors, fusion and tubulation — the situations in which re-seeding was needed in practice. The contour fails on rings below three PSF widths in radius and, at the default integration width, on strongly elongated shapes (a 3:1 ellipse collapsed onto one end, axis ratio 2 was followed; a 2 pixel width follows both), which excludes the smallest early endosomes and the most elongated, tubular intermediates.

On individual, unenlarged endosomes the Rab5-to-Rab7 conversion is sigmoidal and completes within tens of seconds to a few minutes — an order of magnitude faster than published estimates [13, 14, 23] obtained on enlarged endosomes, at minute intervals or as ensemble averages. The digital-switch shape agrees with the bistable Rab5 activation of a reconstituted system [22] and the cut-out switch of network models [21], and the stepwise, cluster-associated Rab5 loss echoes previously described self-organized Rab5 domains on endosomal membranes [21, 37]. The speed varies more than tenfold between endosomes of the same cell line, and the Rab5–Rab7 lag is positive in most events but broadly and asymmetrically distributed, arguing against a fixed coupling and for a variable step — most plausibly recruitment or activation of Mon1–Ccz1 by active Rab5 [15, 19], coordinated with TBC1D18 [20] — whose kinetics differ between endosomes. All measurements were made under transient overexpression (GFP-Rab7a near the endogenous level in MelJuSo cells, Supplementary Fig. 7; Rab5 and regulator levels not determined), so differences between lines may partly reflect expression level. In a minority of events in Supplementary Fig. 5b — and in three of the six resolved Rab7 steps of the spinning-disk cell (Fig. 5) — the Rab7 midpoint preceded the Rab5 midpoint; whether this is genuine ordering, an incomplete trace or a dim Rab7 signal cannot be decided from these numbers.

The placements of other proteins relative to the switch (Fig. 7) rest on few events: only EEA1, Rab14, Rab8 and Rab4 are supported by replicates. EEA1’s departure within seconds of Rab5, as expected for a direct effector [23, 36, 38], validates the measurement; the early departure of Rab14, about 10 min before Rab5 leaves, is the most robust new placement, consistent with a Rab14-dependent step separable from the Rab5/Rab7 pathway [39–41]. That Rab9 and CD63 gain intensity yet are already present on most Rab5 endosomes extends the distinct Rab domains seen on single endosomes [42] into the time domain. These data also suggest one source of the broad, skewed spread of Rab5 τ. No protein was on all Rab5 endosomes (Fig. 7i), so the Rab5 population is a mixture of coats, and a protein that changes in phase with Rab5 loss or Rab7 gain (Fig. 7j) or shifts the speed when overexpressed (Fig. 6a) could set the rate on the subset it occupies. This is a hypothesis; a third color, measuring τ on marker-positive and marker-negative endosomes in the same cell, would test it. The Rab22 results (Fig. 8) agree with the proposed Rab22–Rabex-5–Rab5 cascade [43] and with the fusion-promoting activity of active Rab22 [44–46]; in the single event captured, Rab22 left the endosome about 8 min after Rab5 (Fig. 7c), and GDP-locked Rab22 S12N left the kinetics unchanged while the endosomes shrank during maturation (median diameter ratio 0.74; Fig. 8e). These are overexpression phenotypes; depletion of the proteins or endogenous tagging is the next step before any placement is read as mechanism.

Because the contour is fitted to the summed image and each channel is only measured in the resulting ROI, a third or fourth color adds a measurement, not a change to the algorithm, and would place several proteins on one endosome in a single pass. The same ring geometry applies to macropinosomes [47, 48], phagosomes, lysosomes and autophagosomes; Table 1 gives the acquisition settings they need. The per-frame contour also supports an angular read-out: a follow-up script reports the number, position, lifetime and lateral motion of intensity puncta along the ring (Supplementary Note 1; Supplementary Fig. 8; Supplementary Software 3). With the baseline established, the switch can be perturbed and watched live: depletion of the GEFs and GAPs that drive it, chemical inhibitors or disease mutations such as the Rab7 variants of Charcot–Marie–Tooth type 2B [49] would show, on individual endosomes, whether altered conversion kinetics cause a phenotype or merely accompany it. SACHET turns a qualitative observation, an endosome changing its coat, into a per-endosome time constant, midpoint and lag with a stated precision, operating window and cost in operator effort.

## Methods

### Cell culture, transfection and plasmids

HeLa, MDCK, MelJuSo, A2058 and HFF cells were cultured in Dulbecco’s modified Eagle’s medium (DMEM; BioWhittaker) with 10% FBS (Sigma-Aldrich), 100 U ml⁻¹ penicillin and 100 µg ml⁻¹ streptomycin at 37 °C and 5% CO₂. HeLa, MDCK and A2058 cells were obtained from ATCC, MelJuSo cells from DSMZ, and HFF cells were a gift from Alexander Bershadsky (Mechanobiology Institute, Singapore); lines were not further authenticated. Cultures were confirmed negative for mycoplasma by PCR at regular intervals. Cells were seeded into 35 mm glass-bottom dishes (MatTek) 48 h before imaging and transfected 24 h before imaging with jetOPTIMUS (Polyplus-transfection) according to the manufacturer’s protocol, using 0.5–1.5 µg DNA per construct, the amount being chosen to match the expression levels of the two constructs; at least two independent cell preparations were imaged per condition, although single-event placements in Fig. 7 necessarily come from one preparation.

The MDCK line carrying the invariant chain (Ii, CD74) under an inducible promoter (pMep4) has been described [50–52]; where it was used it was imaged without induction, so that its endosomes were of normal size, as in parental MDCK cells; the one induced session was excluded (Fig. 7 legend). mCherry-Rab5a has been described [53] and EGFP-Rab5a [23, 50]; mApple-Rab5a was from the Bakke laboratory. pEGFP-Rab7a was a gift from Cecilia Bucci (University of Salento) [54]; mApple-Rab7a (plasmid #54945), EGFP-Rab22a (plasmid #49600) and EGFP-Rab4A (plasmid #49434) were purchased from Addgene. mCherry-Rab22a, mCherry-Rab22a Q64L and mCherry-Rab22a S12N were a gift from Subba Rao Gangi Setty; GFP-Rab8 and GFP-Rab11 were a gift from Alexander Bershadsky; GFP-WASH1 (WASHC1) was a gift from Kay O. Schink (University of Oslo). GFP-tagged Rab6, Rab9, Rab14, Rab31, Rab35, EEA1, CD63, LAMP1 and LIMP-2 (SCARB2) were from the plasmid collection of the Bakke laboratory and are available on request. All fluorescent proteins are N-terminal fusions and constructs are named fluorophore–protein accordingly. Because some constructs were available with only one fluorophore, the tag on Rab5 and on the co-expressed protein was swapped between experiments; in all figures Rab5 is displayed in magenta and the co-expressed protein in green irrespective of the tag, and the tags used are stated in the legends.

### Live-cell microscopy

Live-cell imaging was performed in complete DMEM with 10% FBS at 37 °C and 5% CO₂ in a stage-top incubator on one of three systems: (1) a Yokogawa CSU-W1 spinning-disk confocal (Gataca Live-SR module, used in confocal mode) on an Olympus IX83 stand with a Photometrics Prime 95B sCMOS camera and a 60× or 100× Plan Apo oil objective (1.4 NA) for the 2019–2020 recordings (the 2017 recording of Fig. 5 used the same CSU-W1 on a Nikon Ti stand; see below); (2) a Zeiss LSM 880 confocal with a 63×/1.4 NA Plan-Apochromat oil objective and Airyscan detector; (3) the same CSU-W1 system with the Extended Focus module [33]. In extended focus (EF) acquisition the piezo stage steps the focal plane through the cell depth in synchronized micro-steps during a single camera exposure [33], so that each time point is one optical projection that integrates the signal over the swept depth — photometrically comparable to a mean (sum) projection of a densely sampled z-stack — and is acquired about ten times faster than the stack with correspondingly less photobleaching [33]. EF was developed in our laboratory for the endosome tracking described here [33]. Laser lines were 488 nm for GFP/EGFP and 561 nm for mCherry/mApple; exposure was about 100 ms per channel; laser powers were not recorded. The recordings behind each figure were acquired as follows. Fig. 1b, c and Supplementary Movie 1: MDCK cell co-expressing mCherry-Rab5a and EGFP-Rab7a, system 3 (EF), 1.35 s per frame, 500 frames, 1,764 × 1,764 pixels, 108 nm pixels. Fig. 6 and Supplementary Fig. 5: EF projections (system 3) or z-stacks (system 1, analyzed as mean projections) acquired every 10 s for 15–20 min, according to availability; the frame interval, not the microscope, is the parameter that enters the analysis, since the shortest Rab5 τ observed (about 20 s) is sampled at or above the Nyquist rate at 10 s per frame. Fig. 7 and Supplementary Movie 6: two-channel time series with the frame intervals given in Supplementary Table 4 (5.0–20 s; not recorded for the Rab8 and Rab35 series), systems 1 and 3. Fig. 8 and Supplementary Movie 7: z-stacks analyzed as mean projections, frame intervals 4.4 s (Rab22 wild type) and 20 s (Q64L, S12N). Fig. 7i, k and Supplementary Fig. 6: first frames of two-channel recordings; for z-stacks the single best-focus plane was analyzed (Colocalization analysis). Pixel size was 108 nm unless stated otherwise.

### Spinning-disk recording for the real-data validation

The recording of Fig. 5 (Cell2_10sec_10min; 14 May 2017) is of an MDCK cell co-expressing EGFP-Rab5a and mApple-Rab7a, acquired on system 1 (Yokogawa CSU-W1 on a Nikon Ti stand, Live-SR module switched off; MetaMorph 7.10) with a 100× Plan Apo VC 1.4 NA oil objective and a 16-bit sCMOS camera at 110 nm pixels, 488 and 561 nm excitation, 100 ms exposure per channel and plane: 11 z-planes (step 0.453 µm), 61 frames at 10.04 s, 1,024 × 1,024 pixels (113 µm field containing several cells). Analysis was performed on the z-maximum-intensity projection. The cell was tiled into 96 overlapping 11 × 11 µm crops (100 × 100 pixels, 20-pixel overlap), each with a contact sheet of every third frame (Fig. 5a, b); for every conversion seen on a sheet the operator recorded the ring center, the frames at which the ring was clearly Rab5 and clearly Rab7, ring or punctum, and a confidence score (3, textbook; 2, good; 1, possible), and the same event in overlapping tiles was merged by position. In this cell membrane-labeled endosomes of 0.4–0.9 µm radius are resolved as rings, the perinuclear region is dense, and the tracked endosomes move a median 0.1–0.4 µm per 10 s frame with displacements of up to 2.3 µm between consecutive frames (2.9 µm in run M043, a probable switch to a neighboring ring). Each marked event was analyzed in a 101 × 101 pixel crop over the full frame range. The unsupervised SACHET run used automatic re-seeding from the last valid center; Cellpose 3.1 (cyto3 and the fine-tuned model of the crowding benchmark) segmented the summed, percentile-normalized crop at diameter 2R + 2 pixels (R, median supervised contour radius) and followed the mask nearest the previous center. A frame was on target when the unsupervised or Cellpose center lay ≤ 0.5 µm from the supervised SACHET contour centroid of the same frame, and oracle corrections were counted as in the crowding benchmark (Supplementary Methods; run_cellpose_realdata.py).

### SACHET algorithm

SACHET is a JavaScript script for ImageJ/Fiji with a companion contour-fitting plugin; the operator seeds it with a point ROI inside the endosome of interest in the first frame to be analyzed, and the per-frame steps are those of Fig. 1a (Results). Contour failure — typically after a displacement of more than half the ring radius, an out-of-focus neighbor or a fusion event — is corrected by placing a new center in the failing frame and re-running from there. The analysis parameters are the number of radial profiles (20 by default), the radial search length, the integration width of each profile (20 pixels by default), the sub-pixel oversampling factor (10 by default) and the PSF width (Gaussian σ); lengths are given in pixels. They are set at the top of the script and stored with each run; further details are given in the code (Supplementary Software 2). All experimental data and the benchmark of Fig. 3 were analyzed with the default integration width of 20 pixels. At this width each profile is averaged over ±10 pixels perpendicular to the ray, which for rings of 4–6 pixel radius covers much of the lumen: the averaged profile peaks about 1 pixel inside the membrane (the constant radius bias of Table 1, corrected by adding 1 pixel), the contour is insensitive to the exact seed position, and rings of radius ≥ 8 pixels are tracked unstably. An integration width of 2 pixels removes the radius bias (−0.13 ± 0.04 pixels), tracks 8 pixel rings and the 3:1 ellipse of Supplementary Fig. 3, but on the spinning-disk recording of Fig. 5 the unattended track is lost 2.7 times as often (Supplementary Fig. 2; Supplementary Table 6; Supplementary Data 3); the published values were therefore kept at 20 pixels, and 2 pixels is recommended for rings larger than 6 pixels in radius. The PSF width was σ = 1.5 pixels for all experimental data; the radial search length was 10 pixels for the spinning-disk recording of Fig. 5 and, by the default of the script version used, 10 pixels for the confocal recordings of Fig. 1 and Supplementary Fig. 5; benchmark settings are given below. Implementation details, the output table and the correction of the public script are given in Supplementary Methods.

### Arctangent fitting

Membrane intensity traces were fitted with the arctangent model

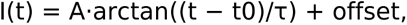

where A is the amplitude swing (negative for a falling channel), t0 the transition midpoint and offset the intensity at the midpoint. With this convention the curve passes from 25% to 75% of its total swing between t0 − τ and t0 + τ, and from 10% to 90% between t0 − 3.1 τ and t0 + 3.1 τ; τ is reported in seconds. Fits were accepted when the transition amplitude exceeded three times the residual standard deviation, R² exceeded 0.5, and the arctangent model fitted better than a monoexponential model (Supplementary Methods). The phase shift between two channels is the difference of their fitted midpoints, Δt0. Choice of model, fitting criterion and preprocessing are given in Supplementary Methods.

### Template-constrained fitting of partially captured transitions

Maturation events are not synchronized with the acquisition window, so within a 15–20 min recording a subset of transitions is captured only partially: the midpoint may precede the start of the recording (left-censored), the transition may still be in progress at its end (right-censored), or the post-transition plateau may not be reached. Free arctangent fits to such traces are poorly constrained, and excluding them would bias the analysis towards fast transitions, since slow events are more likely to be clipped. Only endosomes on which both channels were detectable entered the phase-shift analysis; the fraction of Rab5 endosomes carrying each marker is reported separately (Colocalization analysis). To recover partial events, the complete arctangent fits of the Rab5 loss and Rab7 gain traces of Fig. 1c were normalized to unit amplitude and rescaled by their time constant to give a dimensionless template; partial traces were fitted with this shape held fixed, leaving the midpoint, amplitude and offset free (and the time-scale factor where the observed fraction of the transition allowed it to be identified). Each event was classified as complete, left- or right-censored, and the fraction of the 10–90% transition observed was recorded. When the time-scale factor is free the template fit is mathematically identical to the free arctangent fit, so the two agree exactly on complete events. The sensitivity to truncation was quantified by artificially truncating the complete Fig. 1c traces and refitting: with ≥ 70% of the 10–90% transition observed the midpoint was recovered to within 4 s for Rab5 and 3–21 s for Rab7, whereas with ≤ 50% observed the fit collapsed (midpoint errors of 46–126 s, τ tending to zero). Events with < 70% of the transition observed were excluded. Where more than one event was available for a marker, traces were aligned on the Rab5 midpoint (or on the marker midpoint where Rab5 was the weaker channel), normalized to their mean, binned, and the template was fitted to the pooled curve; 95% confidence intervals on Δt0 were obtained by bootstrapping over events. Alignment removes the jitter of the alignment channel, so its pooled τ is biased low and that of the other channel high; Δt0 is unaffected and is the quantity reported. One exception is stated in the legend of Fig. 7c: the Rab5 trace of the single Rab22 event did not reach its plateau and was fitted with an exponential.

### Synthetic ground-truth benchmark

Synthetic two-channel series were generated in Python (make_benchmark.py; every generating parameter and the frame-1 seed coordinate are listed in manifest.csv). Each frame contains one membrane-labeled endosome modeled as a spherical shell of half-thickness 0.8 pixels around a sub-pixel center at radius R, with luminal intensity 10% of the membrane, convolved with a Gaussian point-spread function of σ = 1.5 pixels (162 nm at 108 nm per pixel) and sampled on a 96 × 96 pixel grid (128 × 128 pixels for R = 20 pixels). Channel 1 falls and channel 2 rises with arctangent kinetics (τ₁ = 60 s, τ₂ = 90 s, midpoints 450 and 570 s, 10% residual floor unless varied) on a background of 100 counts with Poisson noise, 2 counts of read noise and six moving distractor puncta (Gaussian spots of PSF width, random walk 0.8 pixels per frame); the endosome performs a random walk of 0.3 pixels per frame. SNR is the membrane peak above background divided by the standard deviation of the background per pixel. The series are: A, SNR 0.5–30 at R = 6 pixels; B, R = 3–20 pixels at SNR 8, with photon-conserving 2 × 2 and 3 × 3 binned copies; C, τ₁ = 20–240 s with τ₂ = 1.5 τ₁ and a fixed 120 s lag; D, random-walk steps of 1.0 or 2.5 pixels per frame, with or without a drift of 0.4 pixels per frame; E, a rotating ellipse (axis ratio 1.1–1.5) and a radius pulsating by ± 25%; F, exponential photobleaching (900 s), 25 bright distractor puncta, and channel 2 at 30% brightness; G, 300 frames at 1 s with τ = 24 and 42 s and a 13 s lag, i.e. the kinetics of an earlier fit of the Fig. 1c event (the final fit gives τ = 26.7 and 61.1 s and an 8 s lag; the series was not regenerated); H, three replicate realizations at SNR 5; P, one endosome of 0.86 µm radius rendered at 108, 144, 216 and 324 nm pixels with the photon budget conserved. All other series have 100 frames at 10 s. For every frame the true center, radius, axis ratio and the noise-free mean membrane intensity of each channel were stored.

Stacks were seeded and tracked by an operator in Fiji exactly as the experimental data, with the PSF σ set to the rendered value and the radial search length to about 1.4 R + 4 pixels; any manual re-seeding was recorded. From the SACHET output we computed per frame the center error (distance from the fitted contour centroid to the true center), the radius error (median contour radius minus true radius) and the fraction of low-quality fits. A frame was scored as a low-quality fit when the raw 20-point contour had a circularity below 0.6 although the true outline in these series has a circularity of at least 0.9; such frames arise when radial profiles lock onto a wrong intensity maximum (noise, a neighbor or the opposite wall) and are the frames in which the raw 20-point polygon is corrected by the contour closure (Supplementary Methods). Their fraction per run is the fit instability. The criterion requires a known outline and is not applied to experimental data, where a low-circularity contour on a deformed endosome is a measurement, not a flaw (for example M007, Fig. 5d). A run was classed as tracked when the fit instability was at most 10%, as unstable when it was higher or the track had to be re-seeded, and as failed when the centroid was more than 2 pixels off target in most frames or the contour could not be initialized. τ and t0 were obtained from the mean intensity in the tracked membrane ROI with the arctangent fit above and compared with the fit to the noise-free membrane intensity of the same series (ideal tracker; i d e a l _ f i t s in score_comparison.py and the truth fits in score_sachet.py). Scoring is implemented in score_sachet.py; the stacks, ground truth, SACHET output tables and per-run scores (benchmark_master_scores.csv) are distributed with the code.

### Deformation, crowding and generic-tracker benchmarks

Ten deformed shapes (make_deformation_series.py) and 110 crowded stacks (22 conditions × 5 realizations) were rendered with the imaging, noise and kinetic parameters of the round series. Fixed-circle, weighted-centroid, threshold, ring-fit and trackpy baselines were seeded at the frame-1 lumen center and run on the round, deformed and crowded stacks, unattended and, in crowding, with oracle corrections. Shapes, sweeps, neighbor models, scoring criteria and baseline settings are given in Supplementary Methods.

### Comparison with TrackMate and Cellpose

TrackMate 7.14 [27] ran headless in Fiji on the summed channels with the Laplacian-of-Gaussian (LoG) blob detector, which responds to bright spots of a chosen radius, sub-pixel localization and the sparse linear-assignment-problem (LAP) tracker (gap closing within R, two frames; no splitting or merging); the track starting nearest the seed was scored and intensities were read in a ± 2 pixel band at the nominal radius around its center. The detector was tuned on a subset (realization 1 of all 23 crowding conditions, all 8 deformed shapes, 10 round stacks spanning SNR 1.5–15, radius 5–12 pixels and pixel size 108–216 nm) over 24 configurations: LoG radius {R + 2σ_PSF + 1.5 pixels, R + σ_PSF, 0.75 × the former} × quality threshold {Otsu, 0.5 × Otsu} × median filter {off, on} × linking distance {0.5 R, R}. The configuration maximizing the mean over regimes of mean frames on target was kept; configurations within one percentage point were ranked by center error, then by crowded identity switches (Supplementary Data 1). The selected configuration (outer radius, 0.5 × Otsu, median filter on, linking 0.5 R) was then run on all 165 stacks. With the LoG radius equal to the ring radius the detector responds to the membrane rather than the lumen (median offset 0.8 R) and no stack is followed; the difference-of-Gaussians (DoG) detector, TrackMate’s faster approximation of LoG, behaved like LoG. Cellpose 3.1 [29, 55] (cyto3, and cyto3 fine-tuned for 100 epochs on 20 crowded training frames not used for testing) segmented the summed image at diameter 2R + 2 pixels; the mask nearest the previous center was followed. Four intensity read-outs were compared: a ± 2 pixel band at the mask equivalent radius, the filled mask, a 4 pixel inner annulus, and a ± 2 pixel band centered on the membrane peak (the mask boundary moved inward by the median distance between boundary and intensity maximum along 20 rays; median shift ≈ 2.6 pixels, masks ≈ 1.8–1.9 × the true area inside the membrane). The read-out with the lowest median absolute lag error on isolated stacks (round rings and crowding controls, both models pooled) was used throughout (peak band, 4.8 s; filled 7.9 s; annulus 10.6 s; mask-radius band 29.3 s; Supplementary Table 2). Frames on target, center error, identity switches and kinetic errors were scored with the three benchmark scorers; stacks without an acceptable fit (r² < 0.5) are omitted from kinetic metrics. SACHET round-ring values are the operator runs of Fig. 3 (five documented failures counted as 0% on target; three P_snr8 stacks without a SACHET run excluded for all trackers); SACHET deformation and crowding values and all baselines are unattended batch runs. Confidence intervals are percentile bootstraps (10,000 resamples) over stacks, or over whole realizations in crowding (Benchmark_consolidated/).

### SACHET Fiji pipeline

The pipeline of Fig. 2 (SACHET Pipeline v1.0; Supplementary Software 2) comprises five Fiji scripts: TrackMate-based candidate finding with rule-based arctangent filters, unattended SACHET pre-runs, interactive review, event export and interactive SACHET; operator-marked events (Fig. 5a, b) enter at the export step. Presets, candidate criteria, the headless test and the correction of the public SACHET script are described in Supplementary Methods.

### Colocalization analysis

The fraction of Rab5-positive endosomes carrying each protein was scored in two-channel first-frame images of MDCK cells (195 cells, 15 markers) with a custom Python pipeline (Supplementary Software 1) and compared with a per-cell randomized null; segmentation, scoring, exclusions and Manders coefficients are given in Supplementary Methods.

### Statistics

Throughout, n is the number of endosomes (one maturation event each) unless stated otherwise, and all tests are two-sided. For Supplementary Fig. 5, each cell line was imaged in at least two independent experiments with at least 10 cells per line, and events were pooled across cells (n endosomes per line as given in the Supplementary Fig. 5 legend). Rab5 τ, Rab7 τ and the Rab5–Rab7 midpoint lag were compared across cell lines by Kruskal–Wallis test followed by Dunn’s post-hoc test with Holm correction over all pairs (SciPy 1.17, scikit-posthocs 0.17); departure of the τ distributions from normality was assessed with the Shapiro–Wilk test and the sample skewness (SciPy), which is why rank-based tests and medians are used throughout; values are given as median with interquartile range and as mean ± SD, and exact p values are given in the legend. In Fig. 6a groups were compared by Kruskal–Wallis test followed by Dunn’s multiple-comparison test against the Rab5-only or the Rab5/Rab7 group as indicated (GraphPad Prism 9); each group comprises at least 20 cells from at least two independent experiments. In Fig. 8d, e groups were compared by Kruskal–Wallis test followed by Dunn’s post-hoc test over all six pairs with Holm correction (SciPy 1.17, scikit-posthocs 0.17); endosomes were pooled from two independent experiments per group. Confidence intervals on Δt0 in Fig. 7 are event-bootstrap 95% intervals. In Fig. 4 and Supplementary Figs. 3 and 4 medians are over stacks and 95% confidence intervals are percentile bootstraps (10,000 resamples) over stacks or, in crowding, over realizations; tracker comparisons are descriptive and no significance tests were applied. In Fig. 5 and Supplementary Data 2 n is the number of supervised runs (38) on distinct endosomes (37) from one cell; values are medians with ranges and fit parameters are given with standard errors. Significance is indicated as ns, p > 0.05; *, p < 0.05; **, p < 0.01;** *, p < 0.001; ****, p < 0.0001. No statistical method was used to predetermine sample size; no data were excluded other than as stated under Colocalization analysis (Supplementary Methods) and Template-constrained fitting, the 11 of 48 marked spinning-disk endosomes excluded with reasons in Supplementary Data 2, and the synthetic stacks without an acceptable fit noted in the Fig. 4 legend; experiments were not randomized and the operator was not blinded to the condition.

### Literature search

A systematic search with the AI-assisted tool Undermind (report generated 10 October 2025; Supplementary File 1) found no study reporting Rab5-to-Rab7 conversion times for individual mammalian endosomes; query and assessment criteria are given in Supplementary Methods. AI-assisted tools were used for the literature search, for code development, for figure assembly and for language editing; all analyses, results and text were verified by the authors, who take full responsibility for the content.

## Supporting information

Performance of SACHET

## Data availability

The synthetic benchmark stacks with ground truth (round, deformation and crowding series), the SACHET output tables and per-run scores, the tracker-comparison scores (Benchmark_comparison, Benchmark_deformation, Benchmark_crowding, Benchmark_consolidated), the marking sheets, event crops, correction logs and per-event tables of the spinning-disk validation, the source data for all graphs in Figs. 1c, 2–8 and Supplementary Figs. 1–8, the preprocessing test and the Cellpose runs on the spinning-disk events (Source Data file, one sheet per figure), and the per-cell colocalization tables are provided with this paper; the uncropped immunoblots of Supplementary Fig. 7 are available from the corresponding author on request and will be added to the Source Data file. The raw image series underlying Figs. 1 and 5–8 (including the spinning-disk recording Cell2_10sec_10min) and Supplementary Fig. 5 will be deposited in the BioImage Archive and Zenodo during peer review; until then they are available from the corresponding author on request. Source data are provided with this paper.

## Code availability

SACHET (JavaScript for ImageJ/Fiji, with the companion contour-fitting plugin, installation instructions and example data) is freely available at https://github.com/evildna/semi-automated-JavaScript-endosome-tracker-SACHET under the MIT license; the version used here is tagged as release v1.0 and will be archived at Zenodo during peer review. The same repository contains the benchmark generator (make_benchmark.py), the scoring script (score_sachet.py), the arctangent-fitting macro (atan_fitter) and the figure scripts (SACHET_benchmark and Figure folders), and the tracker-comparison scripts (Benchmark_comparison). The colocalization pipeline is provided as Supplementary Software 1 and in the same repository. The SACHET Fiji pipeline (SACHET Pipeline v1.0: scripts SACHET_1–5, the corrected SACHET_Endosome_Tracking_pipeline.js, sachet_run_lib.py, presets.json, user guide and file-format specification) is provided as Supplementary Software 2 (dist/SACHET_Pipeline_v1.0.zip) and in the same repository under the MIT license, together with the Java source of the companion contour-fitting plugin. The public SACHET_Endosome_Tracking.js will be corrected for the intensity-normalization order described in Supplementary Methods before release; SACHET_batch.js (unattended runs), SACHET_batch_reseed.js (automatic re-seeding), make_deformation_series.py, the crowding generator and scorers, and the consolidated comparison scripts (trackmate_sweep.py, run_cellpose_RD.py, cellpose_readouts.py, assemble.py) are in the same repository, as is the ring-puncta script SACHET_Ring_Puncta.js (Supplementary Software 3) and the Cellpose real-data script (run_cellpose_realdata.py).

## Acknowledgements

We thank Subba Rao Gangi Setty (Rab22 constructs), Alexander Bershadsky (HFF cells; Rab8 and Rab11 constructs), Cecilia Bucci (pEGFP-Rab7a) and Kay O. Schink (GFP-WASH1) for reagents, and the NorMIC Oslo imaging platform for access to the microscopes. This work was supported by the Research Council of Norway (INFRASTRUKTUR 208310/F50, NorMIC Oslo; NALMIN-II infrastructure 322607), the Norwegian Cancer Society, and the National Research Foundation and Ministry of Education of Singapore (RCE R-714-105-004-135).

## Author contributions

Conceptualization: X.H., F.M., O.B. Software (SACHET): F.M., X.H. Software (benchmark, comparison, colocalization): X.H. Investigation (imaging): X.H., A.-C.B., D.M., A.V.R., V.M. Formal analysis: X.H., A.-C.B. Resources: X.X., M.P.S., O.B. Writing — original draft: X.H., F.M. Writing — review and editing: all authors. Supervision: O.B., F.M., M.P.S. Funding acquisition: O.B., F.M., M.P.S.

## Ethics declaration

Ethics declaration: not applicable.

## Additional information

Supplementary Information is available for this paper. Correspondence and requests for materials should be addressed to X.H.

